# The genetic architecture of polygenic local adaptation and its role in shaping barriers to gene flow

**DOI:** 10.1101/2023.09.24.559235

**Authors:** Arthur Zwaenepoel, Himani Sachdeva, Christelle Fraïsse

**Affiliations:** CNRS, Univ. Lille, UMR 8198 – Evo-Eco-Paleo, F-59000, Lille, France; Department of Mathematics, University of Vienna, Vienna Austria

**Keywords:** Reproductive isolation, local adaptation, barriers to gene flow, heterogeneous genetic architecture, dominance, haploid selection, life cycles

## Abstract

We consider how the genetic architecture underlying locally adaptive traits determines the strength of a barrier to gene flow in a mainland-island model. Assuming a general life cycle, we derive an expression for the effective migration rate when local adaptation is due to a polygenic trait under directional selection on the island, allowing for arbitrary fitness and dominance effects across loci. We show how the effective migration rate can be combined with classical single-locus diffusion theory to accurately predict multilocus differentiation between the mainland and island at migration-selection-drift equilibrium and determine the migration rate beyond which local adaptation collapses, while accounting for genetic drift and weak linkage. Using our efficient numerical tools, we then present a detailed study of the effects of dominance on barriers to gene flow, showing that when total selection is sufficiently strong, more recessive local adaptation generates stronger barriers to gene flow. We show that details of the haplodiplontic life cycle can be captured using a set of effective parameters, and consider how the relative strength of selection in the two phases affects barriers to gene flow. We then study how heterogeneous genetic architectures of local adaptation affect barriers to gene flow, characterizing adaptive differentiation at migration-selection balance for different distributions of fitness effects. We find that a more heterogeneous genetic architecture generally yields a stronger genome-wide barrier to gene flow and that the detailed genetic architecture underlying locally adaptive traits can have an important effect on observable differentiation when divergence is not too large. Lastly, we study the limits of our approach as loci become more tightly linked, showing that our predictions remain accurate over a large biologically relevant domain.

## Introduction

When a population is subdivided across multiple habitats with different environmental conditions, the extent to which distinct subpopulations can maintain locally beneficial genetic variation depends on the rate of migration between them. Migration between populations that maintain divergently selected alleles can generate migration load (a reduction in mean fitness due to the influx of locally maladaptive alleles) or may lead to loss of local adaptation altogether (so-called *swamping* by gene flow) (e.g. Lenormand (2002)). While local adaptation may be driven by a few conspicuous loci (e.g. adaptive melanism in peppermoths (van’t Hof et al., 2016) or pocket mice (Nachman et al., 2003)), it is believed to typically be polygenic, involving alleles of different effect at many loci across the genome (Pritchard and Di Rienzo, 2010; Le Corre and Kremer, 2012; Westram et al., 2018; Barghi et al., 2020; Bomblies and Peichel, 2022; Stankowski et al., 2023).

When local adaptation is polygenic, migration between populations adapted to different environmental conditions will generate linkage disequilibria (LD), i.e. statistical associations, between selected loci, as locally deleterious alleles will tend to reside in the genomes of individuals with recent migrant ancestry. The rate at which individual locally deleterious alleles are eliminated will be affected by these associations, a phenomenon often referred to as a ‘coupling’ effect (Barton, 1983; Kruuk et al., 1999; Feder et al., 2012; Yeaman, 2015; Sachdeva, 2022). Indeed, sets of loosely linked locally deleterious alleles introduced by migrants will be eliminated jointly in the first few generations after migration at a rate which depends essentially on the relative fitness of migrant individuals and their immediate descendants. These coupling effects will in tassociated with the 1 allele in thuern affect the equilibrium migration load and swamping thresholds (i.e. the migration rate beyond which local adaptation is lost). Neutral variation may also come to be associated with locally selected alleles, so that the latter constitute a ‘barrier’ to neutral gene flow, increasing neutral genetic differentiation (as quantified by *F*_ST_ for instance) beyond the single-locus neutral expectation (Petry, 1983; Bengtsson, 1985).

Barrier effects due to divergent selection at many loci may play an important role in the evolution of reproductive isolation (RI), and hence speciation (Nosil, 2012; Barton, 2020). The colonization of a new habitat will often involve selection on polygenic traits and give rise to a subpopulation that exhibits some divergence from its ancestors (Barton and Etheridge, 2018). Conditional on the initial succesful establishment of such a divergent subpopulation, whether or not speciation ensues depends on whether local adaptation can be maintained in the face of maladaptive gene flow (if any), and on the extent to which the partial RI deriving from local adaptation may promote further divergence and strengthen RI (e.g. through reinforcement, coupling of locally adaptive alleles with intrinsic incompatibilities, or the establishment of additional locally beneficial mutations; Barton and De Cara (2009); Bierne et al. (2011); Butlin and Smadja (2018); Kulmuni et al. (2020)).

Despite mounting evidence that local adaptation is indeed often polygenic, little is known about the underlying genetic details and how these influence the maintenance of adaptive differentiation in the face of maladaptive gene flow (Bomblies and Peichel, 2022; Yeaman and Whitlock, 2011; Yeaman, 2015). How many loci are involved? What are the typical effect sizes? Are divergently selected alleles typically closely linked or spread all over the genome? How non-additive is local adaptation? Relatedly, it is currently unclear how the detailed genetic architecture of local adaptation affects the strength of the resulting barrier to gene flow across the genome. These issues do not only arise in the study of local adaptation *per se*, but are also central to speciation research, as we still have little insight in the genomic architecture underlying isolating mechanisms in nature (Ravinet et al., 2017).

In a recent paper, Sachdeva (2022) showed that, when the loci under selection are unlinked, the effects of LD on equilibrium differentiation at any individual locus in a polygenic system can be well described by classical single-locus population genetic theory, provided that the migration rate *m* is substituted by an *effective* migration rate *m*_*e*_ (Petry, 1983; Bengtsson, 1985; Barton and Bengtsson, 1986; Kobayashi et al., 2008), which captures the multilocus barrier effect, i.e. how gene flow at a focal locus is affected by selection against the associated genetic background. The effective migration rate for a neutral locus can furthermore serve as a quantitative measure of RI, i.e. RI = 1 − *m*_*e*_*/m* (Westram et al., 2022). Crucially, *m*_*e*_ depends itself on the frequencies of divergently selected alleles, giving rise to feedback effects where a small increase in migration rate may cause a sudden collapse of local adaptation (i.e. swamping). In Sachdeva (2022) a detailed study was conducted of the joint effects of drift and LD on swamping thresholds and neutral differentiation in the mainland-island and infinite-island models of population subdivision, assuming a haploid sexual life cycle and divergently selected loci of equal effect.

In this paper, we focus on how the genetic architecture of a locally adaptive additive trait determines the strength of a barrier to gene flow and the conditions for maintaining local adaptation. Extending the theoretical framework of Sachdeva (2022), we derive an expression for the effective migration rate at a neutral locus under a polygenic architecture of local adaptation with arbitrary fitness and dominance effects across loci (referred to as a *heterogeneous* barrier), assuming a population with a haplodiplontic life cycle (which includes haplontic and diplontic life cycles as special cases). We use this *m*_*e*_ to build an approximation for the marginal allele frequency distributions at migration-selection balance in a mainland-island model, allowing us to quantify the extent of adaptive differentiation at equilibrium and swamping thresholds at individual loci influencing the polygenic trait.

After showing the accuracy of our approximations, we use these to address a number of questions concerning the relationship between the genetic architecture of divergently selected traits and the strength of the resulting barrier effect across the genome. In particular, we ask: Are barriers to gene flow stronger when locally adaptive alleles are recessive or dominant? How does selection and migration in both phases of a haplodiplontic life cycle affect a barrier to gene flow? Does a more heterogeneous architecture of local adaptation lead to stronger barriers to gene flow or rather the converse? How does the distribution of fitness effects (DFE) of locally adaptive alleles affect equilibrium differentiation and swamping thresholds at individual loci? How does linkage affect the strength of a barrier to gene flow and swamping thresholds?

## Model and Methods

### Haplodiplontic mainland-island model

Here we outline a mainland-island model for a sexual population which may be subject to selection in both the haploid and diploid phases (fig. 1). We think of this model as a caricature of a bryophyte, pteridophyte, algal or fungal population, but as we shall see below, the model encompasses purely diplontic and haplontic life cycles as well. We assume a regular and synchronous alternation of generations, where an island population of *N* haploids (gametophytes) produces an effectively infinite pool of gametes from which 2*Nk* gametes are sampled that unite randomly to form *Nk* diploid individuals (sporophytes), *k* being the number of diploids per haploid individual. The diploid generation produces in turn an effectively infinite pool of haploid spores through meiosis, of which *N* are drawn to form the next haploid generation. Throughout, we assume that sexes need not be distinguished.

**Fig. 1:**
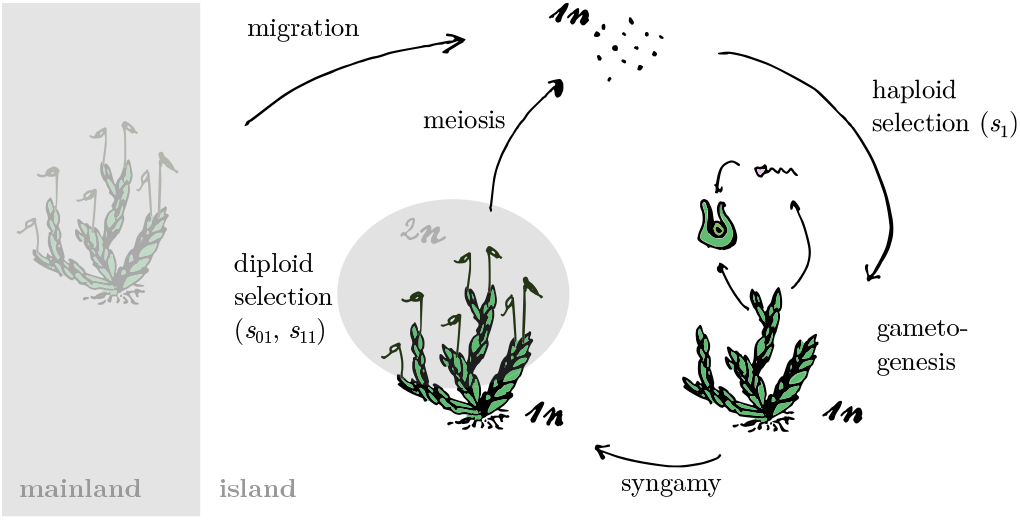
Schematic diagram of the biallelic haplodiplontic mainland-island model (here for a hypothetical bryophyte). Unless stated otherwise, we assume the life cycle to involve migration of haploid (1*n*) spores from the mainland into the island, followed by haploid selection, gametogenesis, random syngamy, diploid (2*n*) selection, and, finally, meiospore formation (meiosis). *s*_1_ is the selection coefficient associated with the `1` allele in the haploid phase, whereas *s*_01_ and *s*_11_ denote the selection coefficients associated with the `01` and `11` genotypes respectively (see main text). We assume a constant number *N* of haploid individuals, with *k* diploid individuals per haploid individual (here *k* = 7). The model is straightforwardly adjusted to allow for diploid or gametic migration.

In each generation, we assume *M* haploid individuals on the island are replaced by haploid individuals from a mainland population, where *M* is Poisson distributed with mean *Nm*. Fitness on the island is determined by *L* biallelic loci which are under divergent selection relative to the mainland. The *L* loci may be linked or unlinked, and we denote the recombination rate between locus *i* and *j* by *r*_*ij*_ . The mainland population is assumed to have a constant, but arbitrary, genetic composition. Unless stated otherwise, we shall assume the mainland to be fixed, at each locus, for the allele which is deleterious on the island. Throughout, we assume a scenario of secondary contact where the locally deleterious allele is initially rare on the island.

Fitness effects are allowed to vary arbitrarily across loci. The wild-type and deleterious alleles (denoted by 0 and 1) at locus *i* have relative fitnesses on the island of 1 and 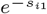 in the haploid phase. The relative fitnesses for the three genotypes in the diploid phase 00, 01 and 11 at locus *i* are 1, 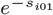, and 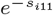. Throughout, we denote the frequency of the wild-type allele (on the island) at locus *i* by *p*_*i*_, and the frequency of the locally deleterious allele by *q*_*i*_ = 1−*p*_*i*_. Fitness is multiplicative across loci, so that, for instance, the log relative fitness of a haploid individual fixed for all the `1` alleles is given by log 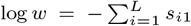. We assume that each haploid (diploid) individual contributes gametes (spores) to the gamete (spore) pool in proportion to its fitness. We assume symmetric mutation at a small constant rate *u* per locus, occurring at meiosis. Individual-based simulations of this model are implemented in a Julia package (Bezanson et al., 2017) available at https://github.com/arzwa/MultilocusIsland.

In the following sections, we build up a theoretical approximation to this model, and validate the approximations by comparing numerical results against individual-based simulations.

### Single-locus theory

We first consider a deterministic model for the allele frequency dynamics at a single locus, ignoring the influence of the other loci as well as genetic drift. As shown in detail in section S2.1, for weak selection and migration, the dynamics of *p* can be described in continuous time by the nonlinear ordinary differential equation (ODE)

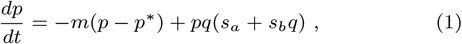

where *p*^*∗*^ is the frequency on the mainland of the allele which is beneficial on the island, *s*_*a*_ = *s*_1_ + *s*_01_ and *s*_*b*_ = *s*_11_ − 2*s*_01_, the latter being a measure of dominance (i.e. the deviation from multiplicative fitnesses (Otto, 2003; Manna et al., 2011)). Usually, *s*_1_, *s*_01_ and *s*_11_ will be assumed to be positive, and *p*^*∗*^ will be assumed to be small, so that selection increases *p*, whereas migration decreases *p*.

When *s*_1_ = 0, we obtain the standard diploid mainland-island model, which is commonly parameterized in terms of a dominance coefficient *h* and selection coefficient *s*, so that *s*_01_ = *sh* and *s*_11_ = *s*. When *s*_1_ ≠ 0 (i.e. there is selection in the haploid phase), we can define an effective selection coefficient *s*_*e*_ = 2*s*_1_ + *s*_11_ and dominance coefficient 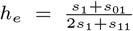, so that, under weak selection, the single-locus dynamics for an arbitrary haplodiplontic life cycle can be described by a diploid model with these effective parameters. Note that *h*_*e*_ is undefined when *s*_*e*_ = 2*s*_1_ + *s*_11_ = 0. We will stick to the more general parameterization in terms of *s*_*a*_ and *s*_*b*_ in our derivations, recalling that these correspond to *s*_*e*_*h*_*e*_ and *s*_*e*_(1 − 2*h*_*e*_) respectively. The equilibria of eq. (1) are analyzed in detail in section S2.1.

When evolutionary forces are sufficiently weak, diffusion theory can be applied to approximate the equilibrium allele frequency distribution at a single locus in a finite population. Note that our model is akin to the standard Wright-Fisher (WF) model with a population size that regularly alternates between *N* and 2*Nk* gene copies. The corresponding effective population size is hence *N*_*e*_ = (*N*^−1^ + (2*Nk*)^−1^)^−1^, twice the harmonic mean of the phase-specific number of gene copies (Hein et al., 2004) (twice because our unit of time is an alternation of generations, not a single generation). The equilibrium allele frequency distribution at a single locus is then given by

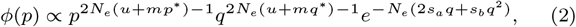

where no closed form expression is known for the normalizing constant. This is essentially Wright’s distribution, generalized to a haplodiplontic life cycle (Wright, 1937).

### Effective migration rate

To make the bridge from single-locus to multilocus theory, we derive an expression for the *gene flow factor* (gff), i.e. the reduction in gene flow at a neutral locus (relative to the ‘raw’ migration rate *m*) due to its being in LD with multiple selected loci (Bengtsson, 1985; Barton and Bengtsson, 1986). As shown formally in Kobayashi et al. (2008), for weak migration, the gff at an unlinked neutral locus equals the expected reproductive value (RV) of migrants in the resident background. At any time, the proportion of individuals with recent migrant ancestry on the island is *O*(*m*), so that the probability of individuals with migrant backgrounds mating with each other to produce, for instance, F2 crosses of the migrant and resident genotypes, is *O*(*m*^2^), and hence negligible for sufficiently weak migration. The descendants of a migrant individual will therefore most likely be F1s and subsequent backcrosses, so that to a good approximation, the RV of a migrant depends only on the relative fitnesses of F1, BC1, BC2, *etc*. individuals.

Let 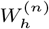 and 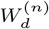 denote the relative fitness of an individual derived from an *n*th generation haploid, respectively diploid, backcross of a migrant with the resident population (i.e. 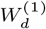 is the relative fitness of an F1 diploid, 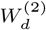 of an offspring from a F1 *×* resident cross (BC1 generation), *etc*.). Assuming migration occurs in the haploid phase before selection, the gff for a neutral locus can be expressed as

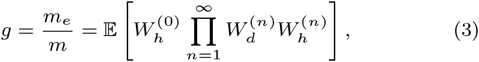

where 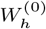 is the relative fitness of the haploid migrant in the resident population (Westram et al., 2022; Barton and Etheridge, 2018; Sachdeva, 2022). Note that this involves an expectation over all possible lines of descent of an initial migrant spore. In practice, *g* is determined only by the first 10 backcross generations or so, as subsequent backcrosses are essentially indistinguishable from residents. While this gff applies strictly only to neutral loci (Kobayashi et al., 2008), it gives a reasonable approximation for the reduction in gene flow at (weakly) selected loci as well.

In order to derive a useful approximate expression for *g*, we shall make two further important assumptions: (1) both the resident and migrant subpopulations, as well as each backcross generation, is in Hardy-Weinberg and linkage equilibrium (HWLE); (2) the expected allele frequency at any locus in any backcross generation is midway between that of the parents (e.g. the mean of the mainland and island allele frequencies for the F1 generation). In reality, due to Mendelian segregation, individuals will not inherit exactly half of the selected alleles of each parent, and this segregation variance will lead to variation within F1s, BC1s, *etc*. on which selection can act. The effects of deviations from the midparent value on the gff are *O*(*s*^2^), so that this last assumption becomes more plausible when local adaptation is due to more and more loci of smaller effect. As derived in Appendix A, in the unlinked case, the approximate gff at locus *j* under these assumptions is

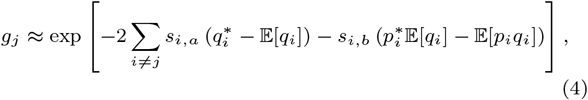

where, again, *s*_*i,a*_ = *s*_*i*1_ + *s*_*i*01_ and *s*_*i,b*_ = *s*_*i*11_ − 2*s*_*i*01_.

It is worth stressing that the gff is a function of the differentiation between the mainland and island population as well as the heterozygosity 𝔼[*pq*] on the island, and that, although we assume migration is sufficiently rare, we do *not* assume that alleles introduced by migrants are rare (i.e. although *m* ≪ 1, *m* ∼ *s*, and hence migrant alleles may have appreciable frequencies). We shall often highlight the dependence of the gff on the allele frequencies and heterozygosities by writing *g*[𝔼[*p*], 𝔼[*pq*]], or *g*[*p*] when allele frequencies are deterministic. Note further that eq. (4) corresponds to 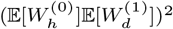, i.e. the product of the relative fitness of a haploid migrant in the haploid resident population and the relative fitness of the diploid F1 in the diploid resident population, squared. Hence, *g* can, in principle, be determined empirically. Equation 4 is straightforwardly adapted to allow for migration in the diploid phase (see Appendix A).

As shown in Appendix B, we can heuristically account for weak linkage by considering the allele frequency dynamics at a neutral locus linked to a single barrier locus, assuming quasi-linkage equilibrium (QLE). The resulting expression for the gff is

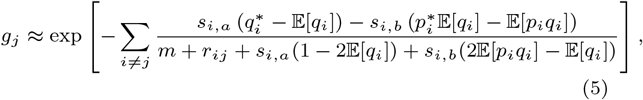

Note that when *r* ≫ *s* ∼ *m* the denominator in the sum in the exponent becomes ≈ *r*_*ij*_, so that eq. (4) appears as a special case of eq. (5) when *r*_*ij*_ = 0.5 for all *i, j*.

### Multilocus dynamics and equilibria

The gff captures the effect of LD among selected loci on the rate of gene flow from the mainland into the island at any individual locus. The key observation is that a certain separation of time scales applies: although selection against migrant *genotypes* can be very strong in the polygenic case (of magnitude *Ls*, roughly), selection at any individual locus is still assumed to be weak, so that, when linkage is weak or absent, LD among selected loci becomes negligible after an evolutionarily short period in which entire sets of alleles are efficiently removed together. Hence, on the longer time scales at which migration-selection balance is attained, the allele frequency at any individual locus should essentially follow the single-locus dynamics, with LD reducing the effective migration rate by a factor equal to the gff (Sachdeva, 2022).

As a consequence, in the deterministic case, we expect that the effects of LD should be well captured by substituting the effective migration rate *m*_*e*_ = *mg* for *m* in eq. (1). Specifically, we get a system of *L* coupled differential equations, where for 1 ≤ *j* ≤ *L*,

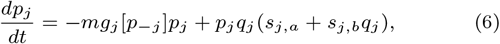

where we assumed the mainland to be fixed for the deleterious allele on the island at all loci. Here we write *g*_*j*_ [*p*_−*j*_] for the gff to highlight the dependence of the gff at locus *j* on the allele frequencies at the other *L* − 1 loci. We study the equilibria of this model by numerically solving for *p* at stationarity (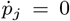, for 1 ≤ *j* ≤ *L*).

We can also plug *m*_*e*_ into the single-locus diffusion approximation to determine the equilibrium allele frequency distribution for each locus on the island (Sachdeva, 2022). Specifically, we can compute moments of the allele frequency distribution at each locus by solving a similar system self-consistently, that is, we solve for the 𝔼[*p*_*j*_] and 𝔼[*p*_*j*_ *q*_*j*_] in

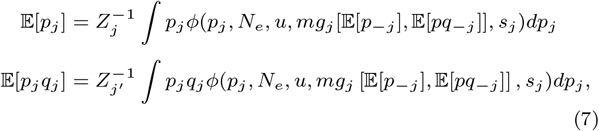

where the *Z*’s are normalizing constants, and 𝔼[*pq*_−*j*_] is the vector of expected heterozygosities at all loci excluding locus *j*. To solve this system of 2*L* nonlinear equations, we use the fixed point iteration outlined in section S2.3. The numerical methods used in this paper are also implemented in the Julia package available at https://github.com/arzwa/MultilocusIsland.

### Realized genetic architecture of divergent selection

For a given DFE of divergent selection, we can use the diffusion approximation to predict the distribution of *s* and *h* (understood as effective selection and dominance coefficients) across the subset of loci that actually contribute to local adaptation for a given rate of migration. We refer to this as the *realized* architecture of local adaptation, and quantify it as the conditional probability density for *s* and *h* at locus *i*, given that a randomly sampled gene copy at locus *i* from the island population is the locally beneficial allele (the event *X*_*i*_ = 1, where *X*_*i*_ is an indicator random variable), i.e.

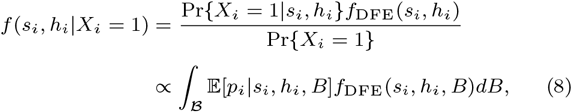

where *f*_DFE_ denotes the joint density of the selection and dominance coefficient across the *L* divergently selected loci, *B* is a shorthand for the selection and dominance coefficients at the *L*− 1 other loci (‘*B*’ for background), and we integrate over the set of all possible such backgrounds *B*. Note that *f*_DFE_ is equivalent to *f* (*s*_*i*_, *h*_*i*_|*X*_*i*_ = 1) in the absence of migration. For a given DFE model, we can characterize this conditional probability density using a Monte Carlo approach by sampling random *L*-locus genetic architectures from the DFE and calculating for each (*s*_*i*_, *h*_*i*_) pair in the barrier the expected beneficial allele frequency 𝔼[*p*_*i*_|*s*_*i*_, *h*_*i*_, *B*] as a weight. The weighted sample will be distributed according to *f* .

## Results

Our main aim is to elucidate how the genetic architecture of polygenic divergent selection determines the strength of a barrier to gene flow, and how, in turn, a polygenic barrier to gene flow affects swamping thresholds at individual loci under divergent selection. We first ignore linkage, and assess the accuracy of our theoretical predictions for the unlinked case by comparing numerical results against individual-based simulations. We next examine the effects of dominance, life cycle details, drift and heterogeneous genetic architectures on the strength of a barrier to gene flow and swamping thresholds. Finally, we consider the effects of linkage between barrier loci for realistic genetic maps.

### Evaluation of the approximation for unlinked loci

We find that substituting *m*_*e*_ = *mg*[𝔼[*p*], 𝔼[*pq*]] for *m* in the single-locus diffusion theory and solving self-consistently for 𝔼[*p*] and 𝔼[*pq*] (see eqs. (4) and (7)) yields remarkably accurate predictions for allele frequencies and swamping thresholds as observed in individual-based simulations (fig. 2). Indeed, even in parameter regimes where the approximation is expected to break down (*Ls* appreciable with *L* small and *s* large, small population size) we obtain good predictions (fig. S1). Not only can we reliably obtain the expected allele frequencies, we also obtain very good predictions for the entire allele frequency distribution (fig. 2, right).

**Fig. 2:**
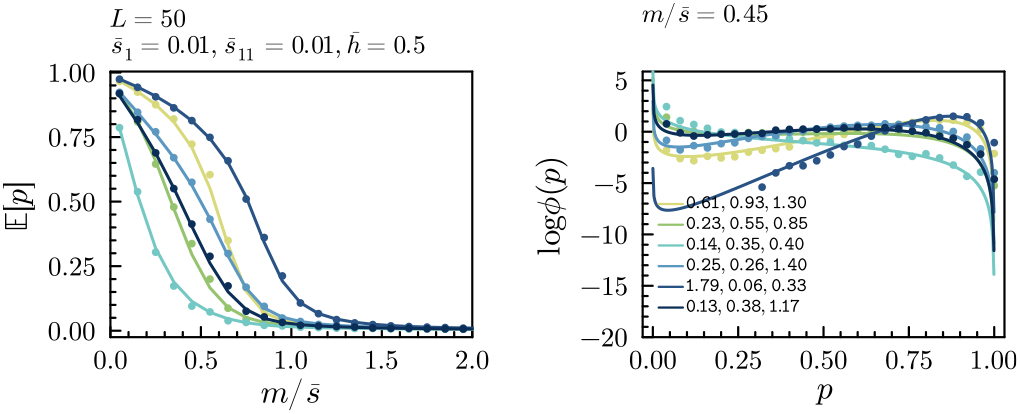
The theoretical approximation for unlinked loci agrees well with individual-based simulations. Left: expected equilibrium frequencies for the locally adaptive allele on the island (𝔼[*p*]) for increasing migration rates for six loci in an unlinked multilocus barrier (*L* = 50). Right: frequency distributions (log_10_ *ϕ*, see eq. (2)) at 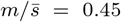 for the same set of loci. Lines show predictions from the multilocus diffusion approximation, whereas dots show results from individual-based simulations (simulating for 200*N* generations, sampling every 10th generation after discarding the first 50*N* generations). We assume a haplodiplontic life cycle with *N* = 550 and *k* = 5, and assume *L* = 50 loci, with selection coefficients in the haploid and diploid phase at each locus sampled from an exponential distribution with mean 0.01, and dominance coefficients for the diploid phase sampled from a Uniform(0, 1) distribution (the values of *s*_1_, *s*_01_ and *s*_11_ for the six highlighted loci are shown in the legend).

### Effect of dominance on barrier strength and swamping thresholds

The effects of dominance on migration-selection balance at a single locus are well known (Haldane (1930); Nagylaki (1975), recapitulated in section S2.2). However, it is less clear how dominance affects the ability to maintain adaptive differentiation and RI for polygenic architectures (but see Harris and Nielsen (2016) for an investigation in the context of selection against introgressed ancestry after an admixture pulse).

To explore the effects of dominance in a polygenic barrier, we write eq. (4) in terms of the effective parameters *s*_*e*_ and *h*_*e*_ (see Methods), and assume all loci have the same fitness effects (a *homogeneous barrier*). Assuming furthermore that the mainland is fixed for the locally deleterious allele on the island, we obtain the simpler expression

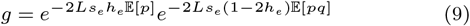

Here, the first factor is just the gff associated with a haploid *L*-locus system with selection coefficients *s*_*e*_*h*_*e*_. The second factor captures the effects of dominance and depends on the heterozygosity 𝔼[*pq*]. Clearly, *h*_*e*_ has opposing effects on both factors. The immediate effect of dominance is therefore that the gff is decreased (barrier strength increased) relative to the additive case (*h*_*e*_ = 1*/*2) whenever invading alleles exhibit a dominant deleterious effect on the island (*h*_*e*_ *>* 1*/*2). Only when heterozygosity (𝔼[*pq*]) becomes appreciable does the second factor contribute to the increase (when *h*_*e*_ *>* 1*/*2) or decrease (when *h*_*e*_ *<* 1*/*2) of the gff.

### Effect of dominance on equilibrium frequencies

To understand better what this means for observable adaptive differentiation, we consider the deterministic multilocus model. From eq. (6), we find that the frequency *p* of the locally beneficial allele at any selected locus in an (*L* + 1)-locus system satisfies at equilibrium

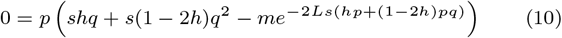

(where we write, for brevity, *s* for *s*_*e*_ and *h* for *h*_*e*_). Assuming *p* ≠ 0, we can divide both sides of eq. (10) by *p* and numerically solve for the equilibrium frequency 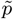. If no such solution exists, the only equilibrium is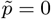.

We find, as expected, that stronger net selection against maladapted genotypes (larger *Ls*) increases the equilibrium frequency of the locally beneficial allele relative to the single-locus prediction, but that the magnitude of this effect depends quite strongly on dominance. When invading alleles are recessive (*h* = 0; fig. 3A), gene flow is not at all impeded when locally deleterious alleles are rare on the island (the gff being near one). This is essentially because, irrespective of how many deleterious alleles a migrant carries, these will not be found in homozygotes as long as migration is sufficiently weak, and hence will not be ‘seen’ by selection. Only once deleterious alleles are segregating at appreciable frequencies on the island, are F1, BC1, *etc*. individuals likely to be homozygous at several loci, thus exposing locally deleterious alleles to selection and reducing the RV of migrants.

**Fig. 3:**
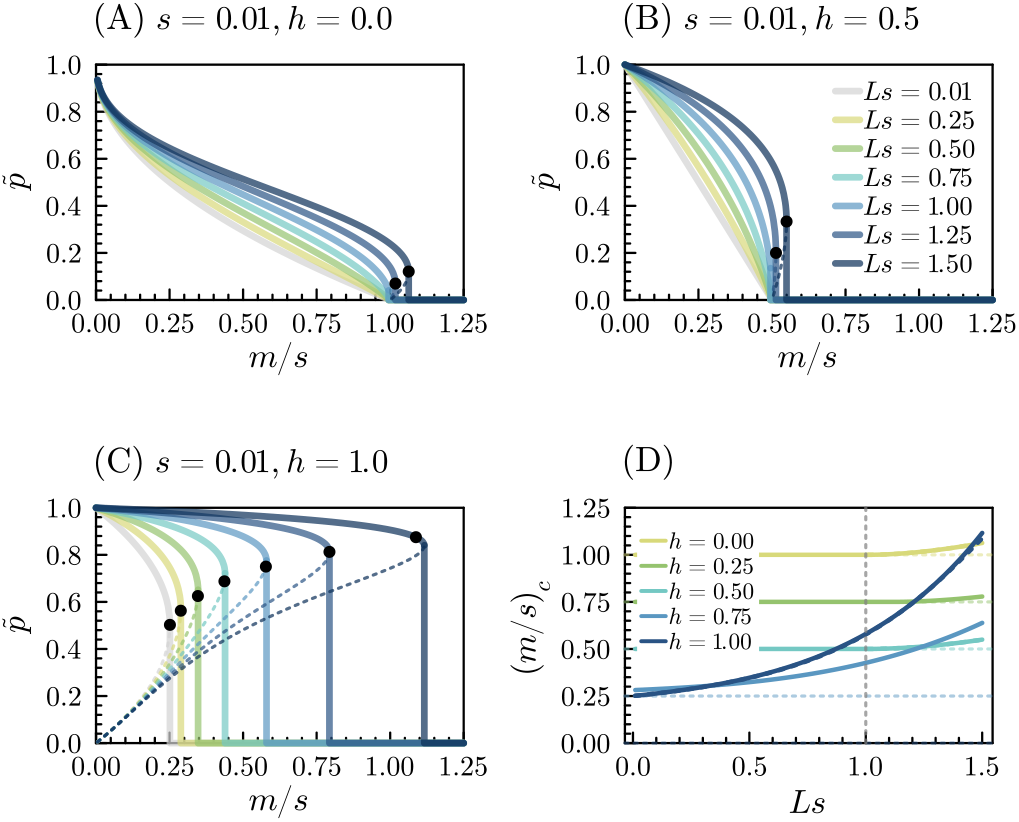
Recessive local adaptation (dominant migrant alleles) leads to stronger barriers to gene flow. Equilibrium frequencies 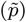 of the locally beneficial alleles in the deterministic model are shown for increasing *Ls* for the case of (A) recessive, (B) additive and (C) dominant migrant alleles. Note that the mainland is fixed for the alternative allele, so that 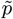 corresponds to the differentiation at equilibrium. The thick lines show the stable equilibria for increasing *m/s*, whereas the dotted lines show unstable equilibria. The black dots mark the critical point beyond which swamping is obtained for any initial condition (in (C) the approximate expression from section S2.4 is used). The results for *Ls* = 0.01 (gray lines) correspond to the single-locus predictions. (D) Swamping thresholds for different degrees of dominance.

The situation is clearly different when migrant alleles are dominant (*h* = 1; fig. 3C), as invading alleles will immediately express their full load in the resident population even in the heterozygous state, causing them to be efficiently eliminated (the gff being at its minimum when migrant alleles are rare). Any increase in the frequency of the deleterious allele on the island will merely increase the expected relative fitness of individuals with migrant ancestry, and hence reduce the barrier effect (increase the gff). We observe a transition between these two qualitatively different types of behaviour at intermediate values of *h*: when *h <* 1*/*3, the barrier strength *increases* with increasing introgression, decreasing the rate of gene flow, until differentiation falls below (3*h* − 1)*/*(4*h* − 2). When *h >* 1*/*3 on the other hand, an increase in the frequency of introgressed alleles always reduces the barrier strength (fig. S3).

### Effect of dominance on swamping thresholds

In the single-locus model, arbitrarily small frequencies of the locally beneficial allele can be maintained at migration-selection balance when *h <* 2*/*3, whereas in the case of *h >* 2*/*3, swamping occurs abruptly if the frequency of locally beneficial alleles falls below 0.5 (section S2.2, gray lines in fig. 3). Sachdeva (2022) showed that such sharp thresholds for swamping also appear in the absence of dominance due to coupling effects. LD both increases the critical migration rate (*m*_*c*_) at which swamping occurs and the minimum level of differentiation that can be maintained before the swamping point is reached (*p*_*c*_). Our results indicate that dominance has a considerable influence on how LD sharpens and displaces swamping thresholds (fig. 3D, section S2.4). For *h <* 2*/*3 (i.e. when local adaptation is not strongly recessive), the critical migration threshold for swamping increases once *Ls* is sufficiently large (as in the additive case), albeit only marginally for *Ls <* 2. This is in sharp contrast with the case where local adaptation is due to strongly recessive alleles (*h >* 2*/*3), where the threshold for swamping increases rapidly with *Ls* (fig. 3D).

Importantly, the critical differentiation (*p*_*c*_) level below which local adaptation collapses is strongly affected by dominance. In the additive case, one can show that sharp thresholds for swamping emerge as soon as *Ls >* 1, in which case *p*_*c*_ = 1 − 1*/Ls*, and hence arbitrary differentiation can be maintained near the critical point depending on *Ls*. For completely dominant local adaptation (*h* = 0), however, *p*_*c*_ increases from 0 to a maximum of 0.5 as *Ls* → ∞, whereas for recessive local adaptation (*h* = 1), *p*_*c*_ increases from 0.5 to 1 as *Ls* grows. This means, in particular, that for moderately strong multilocus selection, *Ls >* 0.75 say, and large population sizes, one would not expect to see locally beneficial recessives at frequencies much below 0.8, compared to 0.5 in the absence of a multilocus barrier effect (fig. 3C).

Figure 3 further highlights that whereas a consideration of the gff in a regime where migrant alleles are rare would suggest that swamping thresholds and equilibria depend roughly on *Lsh*, and not on *Ls* and *h* separately, this intuition really only works well for very small rates of migration (fig. S2).

### Effects of genetic drift

The results for the deterministic model in the previous paragraphs highlight the main effects of dominance on homogeneous polygenic barriers and swamping thresholds. Unsurprisingly, the general consequence of genetic drift is to reduce the barrier effect, hence decreasing the expected differentiation at equilibrium (fig. S4, fig. S5). Swamping thresholds are both decreased and made less sharp by drift, and the detailed behavior depends on the dominance coefficient (fig. S4B).

#### Selection in haplodiplontic life cycles

Figure 2 already showed that we can accurately predict equilibrium allele frequencies for haplodiplontic life cycles with selection in both phases using the diffusion approximation. As the latter depends on *N, k, s*_1_, *s*_01_ and *s*_11_ only through *N*_*e*_*s*_*e*_, *s*_*e*_ = 2*s*_1_ +*s*_11_ and *h*_*e*_ = (*s*_1_ + *s*_01_)*/s*_*e*_, where *N*_*e*_ = (*N*^−1^ + (2*Nk*)^−1^)^−1^ (see eq. (2)), this implies that, at least for weak selection, life cycle details can be accounted for by means of suitable effective parameters (see also fig. S6).

We now consider how the relative strength of selection in the phases affects a polygenic barrier to gene flow. To this end, we parameterize the single-locus model as follows (see also fig. 2)

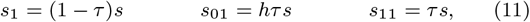

where 0 ≤ *τ* ≤ 1 measures the relative strength of selection in the diploid phase (if one assumes selection to operate with constant intensity throughout the life cycle, this can be interpreted as the relative length of the diploid phase). Similar models have appeared in the study of life cycle modifiers (e.g. Otto (1994); Scott and Rescan (2017)). Figure 4 shows the predicted equilibrium frequencies on the island for homogeneous multilocus barriers where each locus follows eq. (11).

**Fig. 4:**
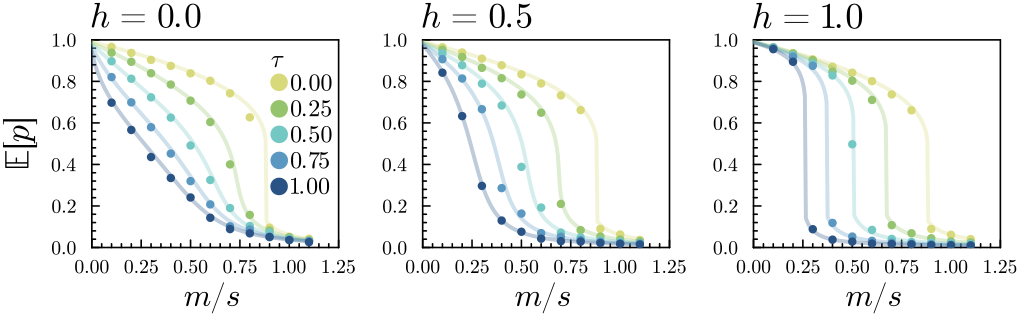
The barrier effect is stronger with increasing haploid selection. The expected differentiation at selected loci is shown for increasing rates of migration and different degrees of dominance in a multilocus model with a general haplodiplontic life cycle. For a given total barrier strength (*Ls*, see main text), we vary the relative strength of selection in the diploid and haploid phase (*τ*, colors) and the degree of dominance in the diploid phase (*h*, columns) (see eq. (11)). *τ* = 1 corresponds to a diplontic life cycle (or at least, absence of selection in the haploid phase), whereas *τ* = 0 corresponds to a haplontic life cycle. Other parameters were: *Ls* = 0.8, *L* = 40, *N*_*e*_*s* = 8, *k* = 5, *u* = *s/*100. The dots show results from individual-based simulations, whereas the lines were computed using the numerical approximation based on diffusion theory.

Under the assumptions of eq. (11), we have *s*_*e*_ = (2 − *τ*)*s* and 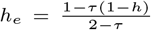 . Hence, increasing the relative strength of selection in the haploid phase (decreasing *τ*) has two important effects. On the one hand, it increases the strength of selection per gene copy. Indeed, in the absence of dominance, the strength of selection per gene copy in a haplontic life cycle (*τ* = 0) is twice that in a diplontic one (*τ* = 1). On the other hand, increasing selection in the haploid phase makes alleles act more additively, that is, as *τ* → 0, we have *h*_*e*_ → 0.5. As we showed that recessive local adaptation can generate stronger barriers to gene flow (fig. 3), this suggests that increasing the strength of haploid selection could weaken a barrier to gene flow as it causes fitness effects to become effectively additive. However, the *joint* effects of *τ* on *s*_*e*_ and *h*_*e*_ are such that, at least for intermediate dominance (0 ≤ *h* ≤ 1), the barrier effect is always stronger with increasing haploid selection (fig. 4). We would therefore expect, all else equal, that barriers to gene flow are stronger in predominantly haploid organisms than in predominantly diploid ones. The relevance of these observations for the evolution and maintenance of haplodiplontic life cycles is however not very clear, as a life cycle modifier need not keep the overall strength of selection constant (Scott and Rescan, 2017).

Lastly, we note that the phase of the life cycle in which migration occurs can influence the maintenance of adaptive differentiation. When there is selection in both phases, and migration occurs in the diploid phase, maladapted diploid migrants are directly exposed to selection on the island, whereas this is not the case when migration occurs in the haploid phase. In the latter case, the first diploid generation exposed to selection is an F1 cross between the mainland and island, so that there is no selection on the homozygous diploid effect. Thus migration in the diploid phase generally leads to stronger barriers to gene flow when selection acts in both phases (fig. S7).

#### Heterogeneous genetic architectures

We now depart from the unrealistic assumption of equal selection and dominance coefficients across the polygenic barrier. In the methods section above, we developed the multilocus theory for general heterogenous genetic architectures (see eq. (4)), where the selection coefficients *s*_1_, *s*_01_ and *s*_11_ can vary arbitrarily across loci, and we already verified that we do indeed obtain accurate predictions in this setting (fig. 2). This allows us to address in more detail a number of questions pertaining to the genetic architecture of local adaptation at migration-selection balance, while accounting for both LD and genetic drift. All results in the following paragraphs apply to a general life cycle with arbitrary selection in both phases, provided that *s* and *h* are interpreted as *effective* selection and dominance coefficients. Given the accuracy of our theoretical approximation (figs. 2 and 4), we restrict ourselves to numerical predictions in the following sections.

### Effect of variation in fitness effects on overall differentiation

We first consider the case with variable selection coefficients across the *L* loci in the barrier, assuming no dominance. Figure 5A shows the average per-locus expected differentiation across the barrier 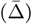 when selection coefficients are sampled from a Gamma distribution (*L* = 100, 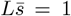, where 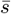 is the average effective selection coefficient across the *L* loci). When migration is weak relative to selection (roughly 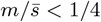), increasing the variance in fitness effects, while keeping 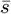 constant, yields on average lower equilibrium differentiation than a homogeneous barrier of strength 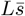, although the barrier strength (as measured by the average gff across the *L* selected loci, 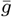) is hardly affected (fig. 5B). By contrast, at higher migration rates, where loci with selection coefficients close to 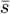 become prone to swamping, heterogeneous architectures tend to yield higher equilibrium differentiation and a stronger barrier effect than a homogeneous one with the same total effect 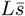 (fig. 5A,B).

**Fig. 5:**
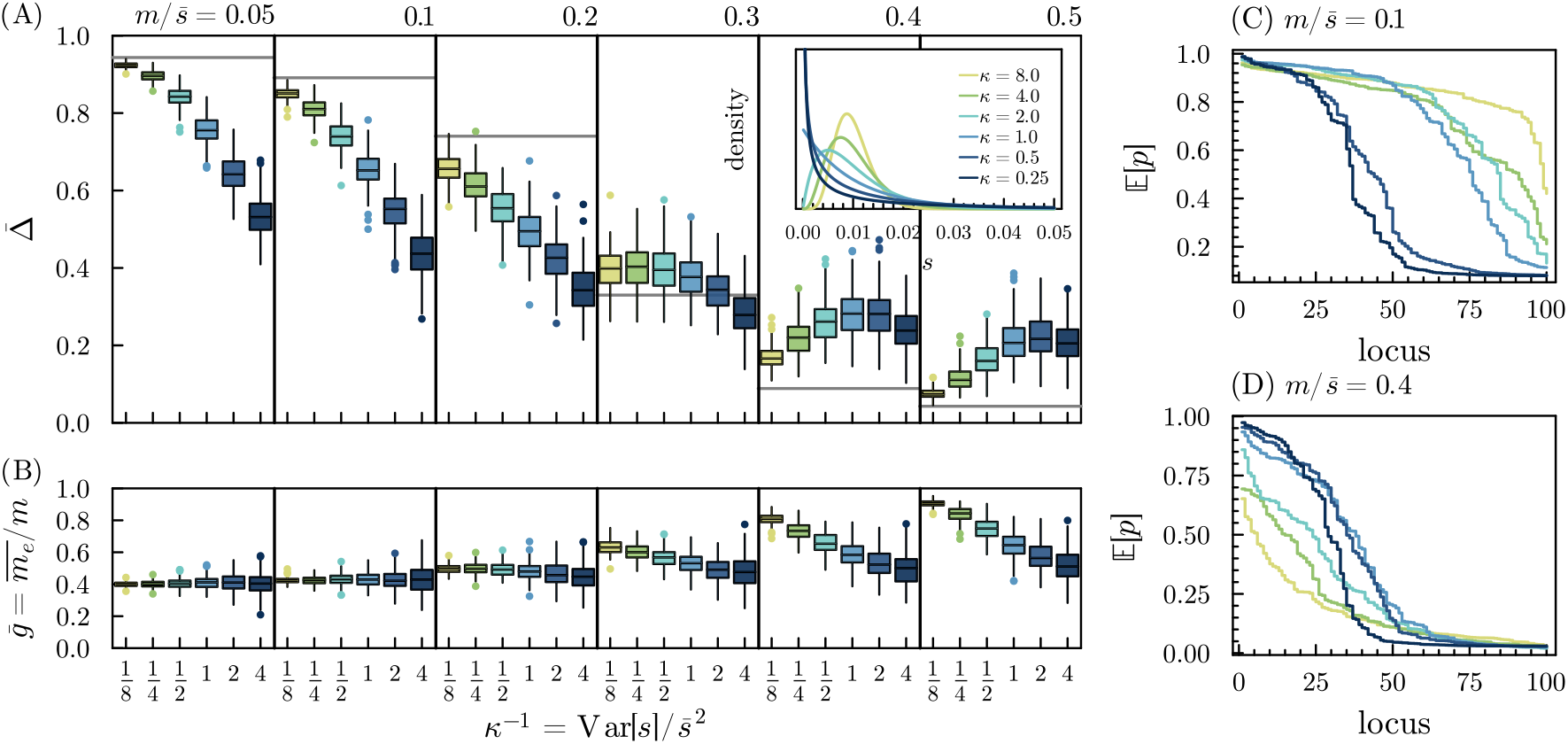
The effects of variation among selection coefficients on polygenic barriers. (A) The boxplots show the mean per-locus differentiation 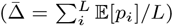 across the *L* = 100 divergently selected loci, for 200 random *L*-locus architectures with no dominance and selection coefficients sampled from a Gamma 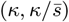 distribution, with 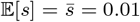 and six different values of *κ* (note that 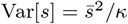 i.e. *κ*^−1^ ∝ Var[*s*]). The solid horizontal line shows the predicted equilibrium differentiation per locus for a homogeneous barrier of strength 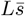 (i.e. the prediction accounting for LD but not for heterogeneity in selection coefficients). (A, inset) Density functions for the six different Gamma distributions used in (A). (B) Average locus-specific gff 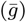 across the *L* divergently selected loci for the same random architectures as in (A). (C,D) Expected beneficial allele frequencies across the barrier in a single simulation replicate for each of the six assumed distributions, sorted by allele frequency, assuming (C) 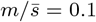 and (D)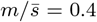 . Colors are as in (A). Other parameters are 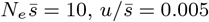 . All results are based on the multilocus diffusion approximation.

One should be careful, however, in the interpretation of 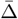 . As shown in fig. 5 (C,D), differentiation across loci in the barrier is often strongly bimodal, especially when Var[*s*] is large, where most loci are either strongly differentiated or not at all, and with rather few loci having 𝔼[*p*] near 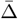 . This implies that empirically, instead of detecting *L* selected loci with an average differentiation of 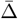, we are more likely to observe about 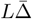 strongly differentiated loci, at least when migration is relatively weak. For low 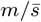, the weaker differentiation observed for more heterogeneous barriers is due to a smaller number of loci effectively contributing to local adaptation, with about half of the locally beneficial alleles swamped at 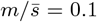 while the other half is strongly differentiated (fig. 5C,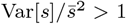) As expected, for stronger migration, the increased differentiation relative to the homogeneous case is driven by a subset of strongly selected loci that resist swamping (fig. 5D).

These results are not significantly affected when there is variation in *h*, at least when *h* and *s* are uncorrelated (fig. S8). The effect of increasing Var[*h*], while keeping the *s*_*i*_ fixed across the barrier, is less dramatic than the effect of heterogeneity in selection coefficients, although we do see systematic decreases or incoefficients, although we do see systematic increases or decreases in equilibrium differentiation depending on whether the migration rate exceeds the swamping threshold for recessives (which are associated with higher equilibrium frequencies) or not (fig. S9). creases in equilibrium differentiation depending on whether the migration rate exceeds the swamping threshold for recessives (which are associated with higher equilibrium frequencies) or not (fig. S9).

### Differentiation and swamping at individual loci in a heterogeneous barrier

Focusing on a single locus embedded within a heterogeneous barrier, we find that, for weak migration, variation in selection coefficients across the barrier has a negligible effect on differentiation at a focal locus with fixed selective effect, whereas (as already shown in fig. 5) it does have a strong effect on average differentiation across the *L* loci (figs. S10 and S11). On the other hand, when migration is strong, a locus with selection coefficient *s* shows on average higher equilibrium differentiation when embedded in a heterogeneous barrier than in a homogeneous one, even when the average differentiation across the barrier is lower in the former (figs. S10 and S11). These results are in line with the observed effect of Var[*s*] on 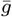 (fig. 5B) and indicate that the presence of a small number of loci of large effect can have a strong impact on the total barrier strength, and hence on the expected differentiation at a focal selected locus when migration is strong (fig. S10). As the distribution of selection coefficients is generally believed to be at least somewhat leptokurtic (note that excess kurtosis ∝ *κ*^−1^ for our Gamma DFE model), we conclude that heterogeneity in selection coefficients can have important consequences for observable differentiation at migration selection-balance that would not be adequately captured by substituting an average selection coefficient in either single-locus or multilocus theory.

Figure 6 highlights how different loci in a heterogeneous barrier are affected differently by the genome-wide barrier effect. For each locus, the expected differentiation at equilibrium is compared against the corresponding single-locus prediction, showing the magnitude of the barrier effect. Selection and dominance coefficients across the *L* loci are assumed to be independently distributed according to a Gamma and Uniform distribution respectively (see section S2.6). As expected, the barrier effect is strongly dependent on both 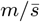 and the total strength of selection 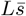 (fig. 6), as well as the strength of genetic drift (fig. S12). Ignoring swamped loci, differentiation is most strongly affected for loci with recessive locally beneficial alleles, whereas the deviation from the single locus prediction for dominant variants is considerably less (figs. 6 and S10). Again we find that when migration is strong, increased heterogeneity of selection coefficients leads to stronger barrier effects, where an appreciable proportion of alleles are protected from swamping due to association with a few strongly selected barrier loci (figs. S13 and S14).

**Fig. 6:**
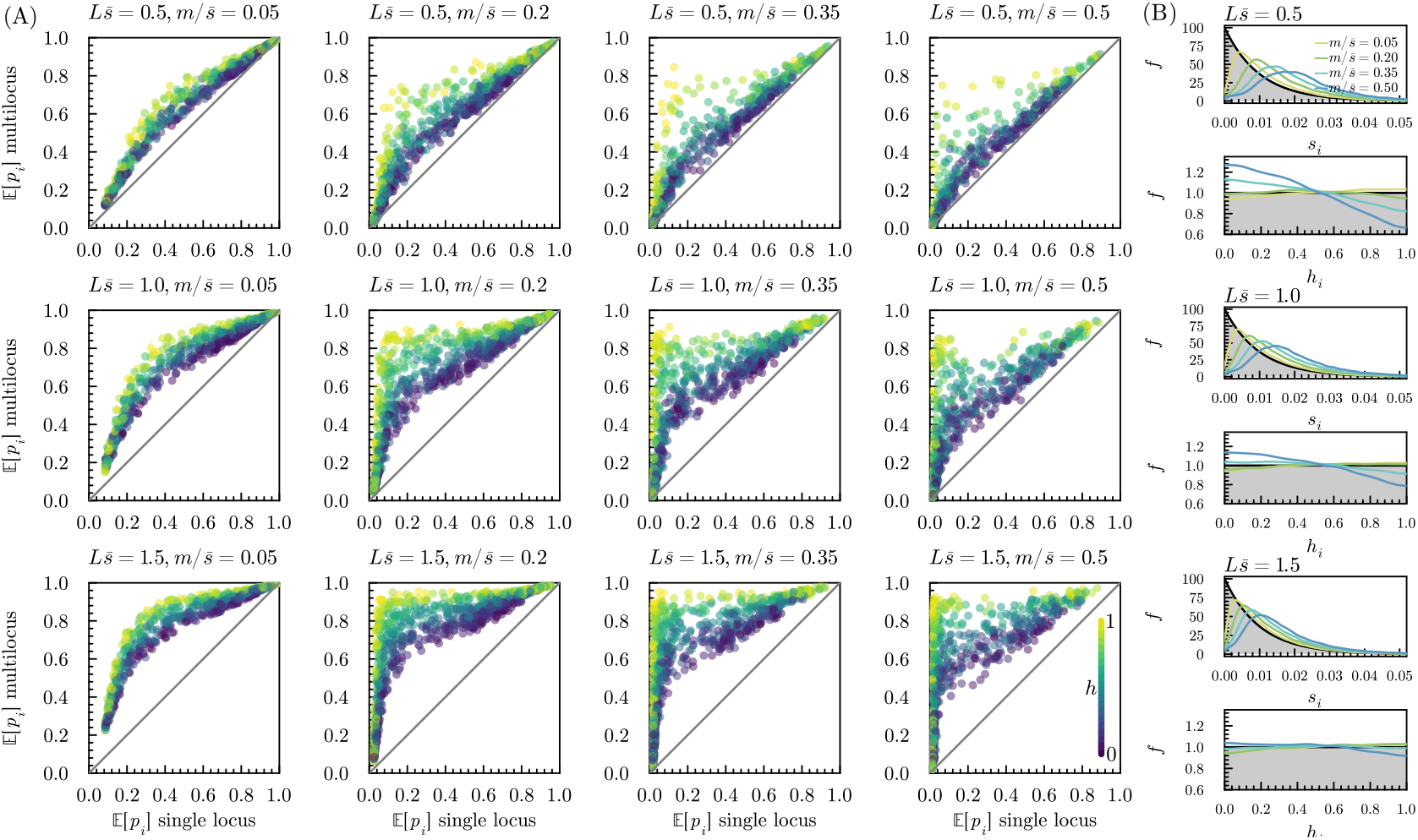
Characterizing barrier effects for heterogeneous genetic architectures of local adaptation. (A) Deviation of predicted allele frequencies for loci in heterogeneous polygenic barriers when accounting for LD (i.e. predictions using the multilocus theory, *y*-axis) from single-locus predictions that neglect LD (*x*-axis). The rows show results for different total strengths of selection 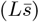, whereas the columns show results for increasing rates of migration relative to selection 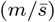 . We assume the *s*_*i*_ to be exponentially distributed with mean 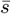 and dominance coefficients are sampled uniformly from the [0, 1] interval. Each dot is associated with a single locus in an *L*-locus barrier, and is colored according to its dominance coefficient (yellow for locally beneficial recessives (*h* = 1), purple for dominants (*h* = 0)). Each plot shows results for 1000 such loci, subsampled from a total of 150000*/L* randomly sampled *L*-locus architectures. (B) Monte Carlo approximation to the marginal distribution of the selection and dominance coefficient conditional on observing a divergent allele on the island (i.e. *f* (*s*_*i*_|*X*_*i*_ = 1) and *f* (*h*_*i*_|*X*_*i*_ = 1), see eq. (8)). The distribution graphed in gray shows *f*_DFE_, i.e. the marginal distribution of the selection and dominance coefficient for a random locus in the *L*-locus barrier in the absence of migration. We assumed 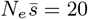 and 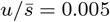for all results.

### The realized architecture of local adaptation

Although the above results indicate that, when 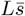 is appreciable, the barrier effect is strongest for recessive alleles (fig. 6), this does not imply that these contribute disproportionally to polygenic local adaptation. Although strongly selected recessives will tend to show strong differentiation, weakly selected recessive alleles will be more prone to swamping than partially dominant ones (e.g. fig. S10). One way to quantify how these two phenomena interact to yield the *realized* genetic architecture of divergent adaptation (related to the concept of *adaptive architecture*, as defined in Barghi et al. (2020)) is by considering the conditional probability density for the selection and dominance coefficient at a locus, given that a locally adaptive allele is observed on the island at that locus (see Methods, eq. (8)). Figure 6B shows approximations to the marginal distributions *f* (*s*_*i*_|*X*_*i*_ = 1) and *f* (*h*_*i*_|*X*_*i*_ = 1) obtained in this way for the heterogeneous barrier model assumed in the preceding section.

As expected, we find that as migration rates go up (colors in fig. 6B), the distribution of selection coefficients in the barrier at migration-selection balance shifts towards higher values of *s*. This effect becomes weaker with increasing 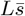, which increases the extent by which small-effect alleles are protected from swamping. Notably, recessives contribute *less* to adaptation than dominants when migration is sufficiently strong (note the blue curves in fig. 6B, corresponding to 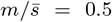), despite the fact that, conditional on no swamping, equilibrium frequencies of recessives are most affected by LD (fig. 6A). This is most notable when 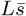 is not large (top row in fig. 6). When 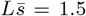 for instance, the depression in the conditional density at *h* = 1 becomes very slight even for relatively large migration rates (bottom row in fig. 6B). We observe a similar shift in the distribution of dominance coefficients when *h* is Beta distributed with mean 2*/*3 instead of uniformly on the unit interval (fig. S15).

Interestingly, we observe a non-zero correlation between *s* and *h* among loci that contribute to local adaptation, even when no such correlation exists across the *L* divergently selected loci. At equilibrium, observed variants of relatively large effect are more likely to act recessively than variants of small effect (figs. 7 and S18). The correlation is negligible for small migration rates, but as the strength of migration increases so that swamping effects become relevant, the correlation coefficient can become as large as 0.25, depending on 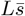.

**Fig. 7:**
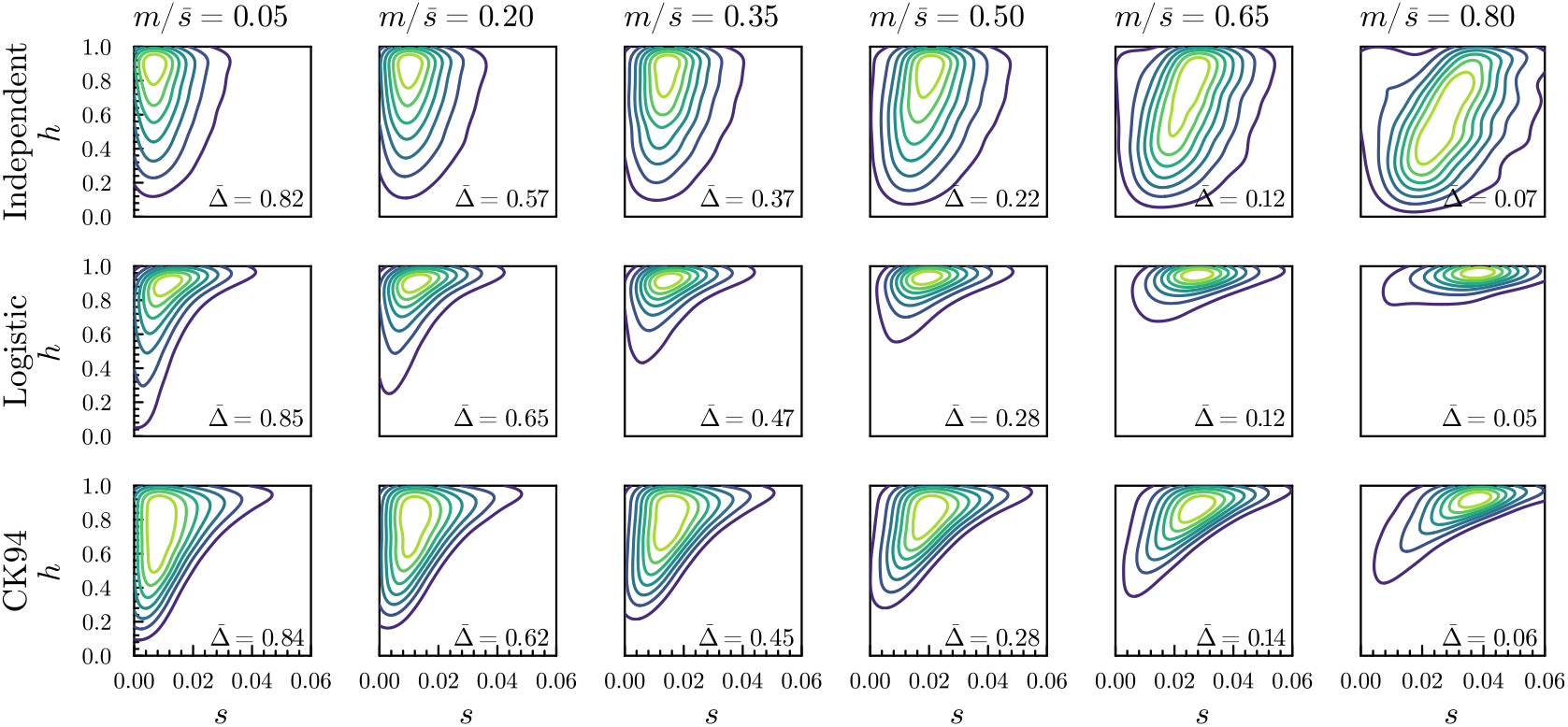
The distribution of fitness effects (DFE) affects the realized genetic architecture of local adaptation at migration-selection balance. Contour plots for the joint density of *h* and *s* conditional on observing a divergent allele on the island (see eq. (8)) are shown for the three DFE models (rows) for increasing rates of migration (columns). Values of 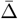 in the lower right corner denote the mean expected differentiation per locus. We assume 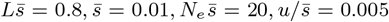 and exponentially distributed selection coefficients, and parameterize the DFE models so that 𝔼[*h*] = 2*/*3, assuming *α* = 2, *β* = 1 for the independent model, *a* = 7.2, *b* = 1.2, *σ* = 1 for the logistic model and *K* = 50 for the CK94 model (see section S2.6 for details on the different DFE models considered here). The densities are approximated using a Monte Carlo approach, simulating 500 replicate *L* locus genetic architectures from the assumed DFE model, determining the equilibrium allele frequencies for each replicate, and fitting a kernel density estimate to the sample so obtained.

The simple DFE model assumed above where selection and dominance coefficients are independent is almost certainly inadequate. Theoretical and empirical work has indicated that, on the one hand, a correlation between selective effect and degree of dominance can be expected in standing genetic variation, with segregating deleterious alleles more likely to be recessive if they have large *s* (Caballero and Keightley, 1994; Zhang et al., 2004; Agrawal and Whitlock, 2011). On the other hand, it is well appreciated that during adaptation, dominant beneficial alleles have higher establishment probabilities than (partially) recessive ones with the same homozygous effect (Haldane’s sieve, Haldane (1927); Turner (1981)). These two aspects interact when adaptation is from standing variation: while more recessive beneficial alleles are less likely to fix than dominant alleles, they are more likely to be segregating at appreciable frequencies when the population faces an adaptive challenge (Orr and Betancourt (2001), but see Muralidhar and Veller (2022)).

To examine how the realized genetic architecture of local adaptation depends on the correlation between *s* and *h* among divergently selected loci, we consider two alternative, admittedly *ad hoc*, DFE models, outlined in section S2.6. Both models assume Gamma distributed selection coefficients and incorporate a positive correlation between *s* and *h*, so that alleles of large effect tend to be more recessive (recall once more that *h* in our case is the dominance coefficient of the invading allele, so *h* = 1 corresponds to recessive local adaptation). We keep the average dominance coefficient fixed to 2*/*3 for each model. In contrast with the independent model, we find that for the model in which *s* and *h* are positively correlated, recessives are typically more likely to contribute to the realized differentiation at equilibrium (figs. 7, S15 to S17 and S19 to S21). When we make the opposite assumption that locally beneficial alleles tend to be dominant (corresponding, for instance, to the case with a strong Haldane’s sieve effect during adaptation), we find a somewhat less dramatic shift in the joint density as migration rates go up (fig. S22). The distribution also shifts towards higher selection coefficients, but somewhat less so than in the model with the opposite correlation. Swamping of partially recessive alleles of small effect further shifts the distribution towards smaller values of *h*. These examples show how correlations between *s* and *h* among the loci under divergent selection can have a rather important influence on the realized genetic architecture at migration-selection balance (i.e. on which loci actually contribute to adaptive differentiation), driving up the relative contribution of recessives in one case but not in the other.

#### The effects of linkage

So far, we have ignored physical linkage of the *L* loci under selection. In realistic genomes, however, it is highly unlikely that there are more than a dozen truly unlinked loci. The importance of linkage will depend on the total map length, the number of chromosomes, and the number and location of the selected loci. For organisms with a small number of chromosomes (e.g. *Drosophila*), linkage may have important consequences, whereas with larger chromosome numbers (for instance, in humans), most of the barrier effect may be due to unlinked loci.

The overall effect of linkage should be to increase the strength of a barrier to gene flow, as linkage increases the variance in introgressed ancestry among a migrant’s descendants, yielding more efficient purging of sets of introgressing maladaptive alleles (Barton, 1983; Veller et al., 2023). However, it is less clear what happens in finite populations and close to swamping thresholds, where linked combinations of adaptive and deleterious alleles may persist due to Hill-Robertson interference. We expect that when recombination is strong relative to selection, i.e. *r*_*ij*_ ≫ *s*_*i*_ for all *j*, the basic separation of timescales between the breakdown of LD and the establishment of migration-selection equilibrium at individual loci still applies, and eq. (5) should yield reasonably accurate predictions. However, associations between tightly linked loci, for which *r* ∼ *s*, will be broken down by recombination at rates comparable to or slower than their elimination by selection, so that the strength of coupling is increased (Barton, 1983; Kruuk et al., 1999), and the approach based on assuming QLE (see Appendix B) becomes inappropriate.

These predictions are verified in fig. 8. As the strength of recombination relative to selection (*r/s*) decreases, the barrier strength increases. We find that for *r/s* ≥ 4, equilibrium frequencies and swamping thresholds appear to be accurately predicted using single-locus diffusion theory with the approximate gff given by eq. (5), although a slight but systematic overprediction of the equilibrium frequencies is apparent. As expected, when the strength of recombination decreases further (*r/s <* 4), the approximation breaks down.

**Fig. 8:**
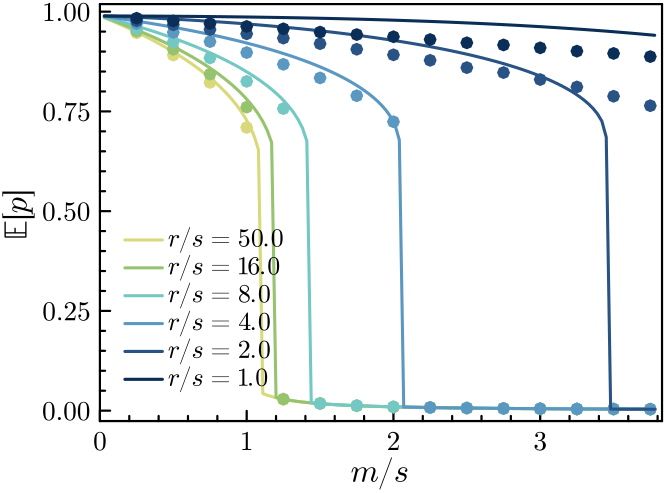
Effects of linkage on barrier strength and on the accuracy of the predictions based on the adjusted effective migration rate (eq. (5)). We assume *L* = 100 loci lying on a single chromosome with recombination rate between adjacent loci equal to *r*, and assume haploid selection with *Ls* = 1, *N*_*e*_*s* = 8 and *u/s* = 100. *r/s* = 50 corresponds to the unlinked case. The map lengths corresponding to *r/s* = 16, 8, 4, 2 and 1 are approximately 1900, 860, 410, 200 and 100 cM respectively. The dots show results (expected frequency of the locally beneficial allele) from individual-based simulations, whereas the lines show predictions based on diffusion theory with the effective migration rate calculated as in eq. (5).

We now consider what this means for realistic genetic maps. In fig. 9 (see also fig. S23), we compare our numerical predictions against individual-based simulations for *L* = 100 equal-effect loci, uniformly distributed along the human and *Drosophila* genomes (i.e. each selected locus is at a randomly sampled position in the genome), assuming *Ls* = 1 and *N*_*e*_*s* = 10. We obtain very good predictions for both equilibrium frequencies and swamping thresholds for the human genetic map, where tight linkage is rare (99% of the *r*_*ij*_ */s* values exceed 10). For the *Drosophila* genetic map (where 92% of the *r*_*ij*_ */s* values exceed 10), we obtain fair predictions for equilibrium differentiation when the migration rate is not too high. However, we tend to predict swamping at lower *m/s* than observed in individual-based simulations, in line with our observations in fig. 8.

**Fig. 9:**
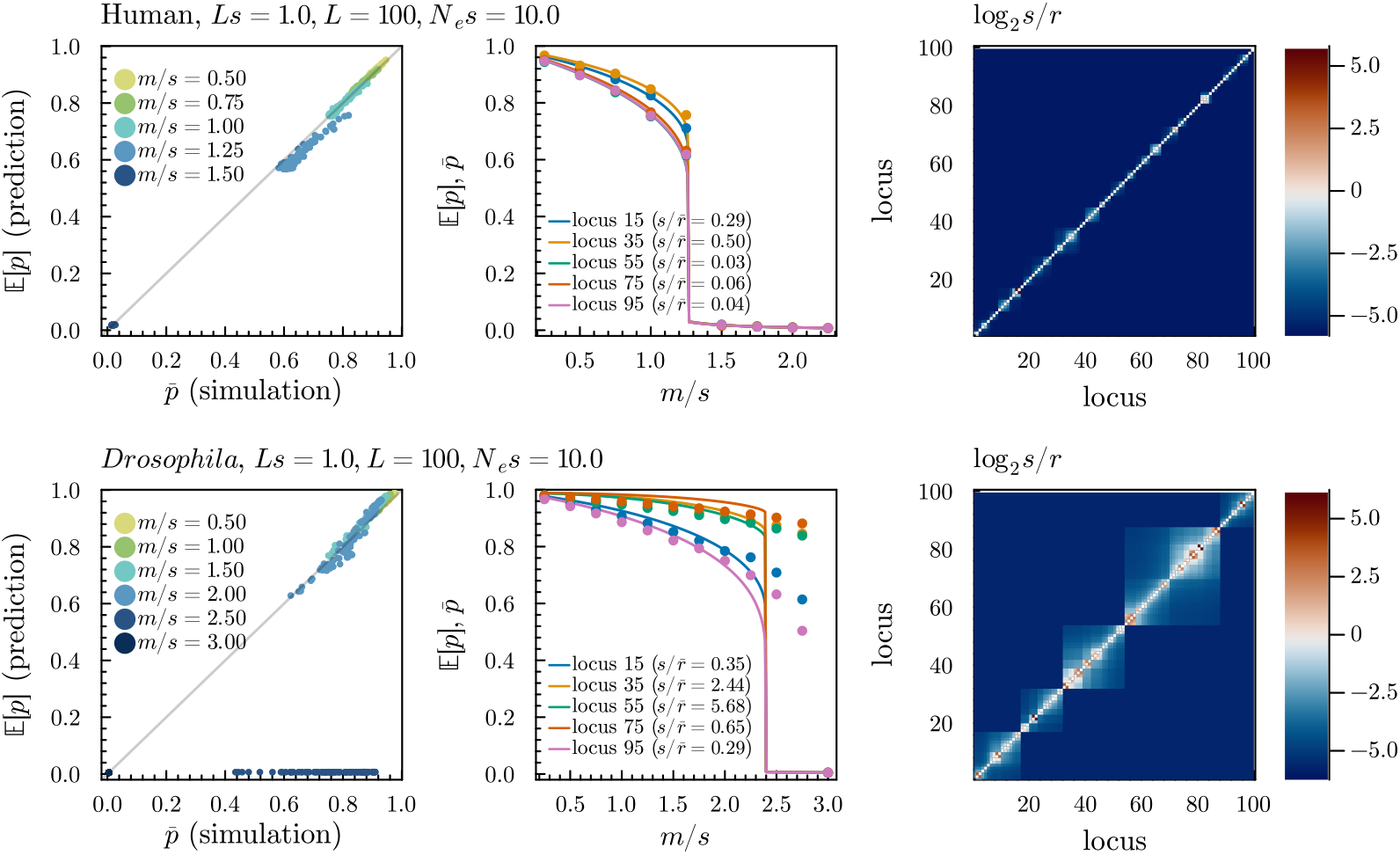
Effects of linkage on the equilibrium frequencies of locally adaptive alleles for *L* = 100 loci randomly scattered on the Human and *Drosophila* genomes. From left to right: scatter plot of the predicted equilibrium frequencies of the locally beneficial allele (𝔼[*p*]) at the 100 loci versus those observed in individual-based simulations 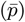; observed (dots) and predicted (lines) allele frequencies for five loci for increasing migration rate; strength of selection relative to recombination across the genome (log_2_ *s/r*), i.e. for each pair of loci *i* and *j*, log_2_ *s/r*_*ij*_ is shown. We assume haploid selection with *Ls* = 1, *N*_*e*_*s* = 10, *u* = *s/*100.

These results highlight that, for similar genetic architectures and population genetic parameters, divergent local adaptation is expected to lead to a much stronger barrier to gene flow (and hence stronger RI) in organisms with a *Drosophila*-like recombination landscape than in organisms that resemble humans in this regard (fig. 9). Prediction accuracy is not noticeably different for heterogeneous barriers (fig. S24); nor does it depend in any obvious way on effective selection and dominance coefficients (fig. S25). This suggests that the effects of linkage are largely orthogonal to the effects of dominance and barrier heterogeneity studied above, at least in the absence of epistasis.

## Discussion

### LD and polygenic migration-selection balance

We study genetic barriers to gene flow comprising an arbitrary number of unlinked or weakly linked divergently selected loci with heterogeneous selective and dominance effects, assuming a mainland-island model and a general haplodiplontic life cycle. We derive expressions for the effective migration rate at any given locus in terms of the divergence levels at all other loci, and then use this together with classical single-locus theory to self-consistently predict divergence at all selected loci. Importantly, this allows us to study the effects of coupling among barrier loci without assuming that locally deleterious alleles are somehow rare, enabling us to study swamping by gene flow in the polygenic setting.

Our results show how the maintenance of adaptive differentiation in the face of gene flow depends jointly on the extent of LD, drift, dominance and variation in selective effects across loci. The general success of the approach indicates two important features of polygenic migration-selection balance. Firstly, it suggests that the ‘separation of time scales’ argument that is at the root of the approach indeed works, and generalizes well beyond the case of haploid equal-effect loci to more realistic architectures with dominance, weak linkage and a haplodiplontic life cycle. Unless loci are tightly linked (i.e. *r/s <* 4), strong selection against multilocus genotypes occurs only in the first few generations after a migrant arrives, and the long term fate of a migrant allele is unaffected by LD conditional on having survived these initial generations. As a consequence, the effects of LD are well described by the usual single locus dynamics, but with a reduced migration rate. Secondly, it indicates that our rather crude approximation to the expected reproductive value of a migrant individual on the island (which assumes HWLE within the island population, that migrants only cross with residents, and that in each such cross the proportion of migrant alleles is exactly halved) is an adequate estimator of the gff.

### The effect of dominance on a polygenic barrier to gene flow

Our analyses for homogeneous genetic architectures indicate that, when there is selection in the diploid phase, dominance can have a considerable impact on the expected level of adaptive differentiation and swamping thresholds. In the multilocus setting, depending on the total strength of divergent selection (*Ls*), partially recessive variants may lead to strongly increased swamping thresholds and produce a much stronger barrier to gene flow than dominant variants with the same homozygous effect (fig. 3), in sharp contrast with the single-locus case (Haldane, 1930). The reason for this is that for recessive local adaptation, the dominant invading alleles are immediately exposed to selection, whereas for dominant local adaptation, the recessive invading alleles can introgress easily as long as the frequency of locally deleterious alleles is low. We find that dominance has a strong effect on the feedback between the level of differentiation and the strength of selection against migrants (which leads to sharp swamping thresholds).

It should be emphasized, however, that all our results assume a mainland-island model of migration and a scenario of secondary contact. The effects of dominance may be more subtle in models with multiple demes and more symmetric patterns of migration, in which case assumptions on environmental dependence of dominance may become important (e.g. Bürger (2013)).

### Life cycle details

We developed our theory for a fairly general haplodiplontic life cycle, and showed that a model with selection in both the haploid and diploid phase can be brought into correspondence with a strictly diploid model through a set of effective parameters. Increasing the relative strength of selection in the haploid phase, for a fixed total strength of selection (*Ls*), yields stronger *effective* selection (larger *s*_*e*_) at a selected locus, while making the fitness effect more additive (i.e. *h*_*e*_ moves towards 1*/*2). One might therefore expect that local adaptation would lead to stronger barriers to gene flow in predominantly haploid species, but it is hard to make more refined predictions at such a general level. This relates to recent work on the strength of barriers to gene flow on sex chromosomes (Fraïsse and Sachdeva, 2021) and in arrhenotokous species (Bendall et al., 2022), where increased exposure to haploid selection generally leads to stronger barrier effects.

Importantly, similar considerations apply *within* the genome of haplodiplontic (and even diplontic) species, as there can be considerable variation in the relative expression levels in the two phases across genes within a genome (e.g. Szövényi et al. (2011); Cervantes et al. (2023)), so that it is likely that the relative strength of selection in the phases varies across the genome (Immler and Otto, 2018). It therefore seems plausible that genes whose expression is biased towards the haploid phase may contribute more to local adaptation, similar to what has been described for sex chromosomes versus autosomes (Lasne et al., 2017), although this would of course depend on the relative extent of divergent selection in both phases.

### Heterogeneous architectures of polygenic barriers

When migration is not strong relative to the average strength of selection per locus, increased variation of *s* in the DFE underlying locally adaptive traits gives rise to lower overall adaptive differentiation at equilibrium, while generating a barrier to gene flow of similar strength (fig. 5). On the other hand, for high rates of migration, more heterogeneous architectures generate a stronger barrier to gene flow due to the presence of a larger number of strongly selected loci that resist swamping. The presence of a few loci of relatively large effect can cause a substantial reduction in gene flow even at unlinked loci, raising neutral differentiation across the genome. As a result, it may become harder to map loci underlying adaptation when the genetic architecture becomes more heterogeneous. At the same time, our results also emphasize that, despite appreciable genome-wide coupling effects, considerable variation in equilibrium differentiation across non-swamped loci remains, depending on the locus-specific fitness effects (fig. 6).

### The genetic architecture of local adaptation at equilibrium

When there is appreciable gene flow, only a subset of the divergently selected loci that underlie a locally adaptive trait will actually exhibit substantial differentiation at migration-selection balance, and the DFE of these loci need not be representative for the DFE associated with all loci underlying the trait (that is, the genetic architecture of local adaptation may differ to greater or lesser extent from the genetic architecture of locally adaptive traits (Yeaman and Whitlock, 2011)).

We find that when selection is fairly weak (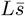 is small), the subset of divergently selected loci that exhibit significant differentiation at migration-selection balance constitutes a more biased subset than when 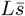 is large (fig. 6). This has implications for our ability to map locally adaptive loci: when 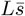 is large, RI is effectively complete, and genome-wide differentiation will be high, so that mapping locally adaptive loci becomes essentially impossible. When 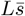 is small, adaptive loci will be more easily detected, but they will constitute a more biased subset of the loci that contribute to variation in locally adaptive traits. Similarly, the DFE at equilibrium shifts more and more to larger selection coefficients when migration increases (fig. 6), whereas the effect on the distribution of dominance coefficients depends on the correlation between *s* and *h* in the DFE underlying locally adaptative traits (figs. 6 and S15 to S17).

Importantly, the maintenance of polygenic migration-selection equilibrium itself *generates* correlations between selection and dominance coefficients. Such correlations are often discussed in the context of adaptation, with different factors influencing the relationship between the homozygous effect and dominance deviation in the DFE of new mutations (Agrawal and Whitlock, 2011; Manna et al., 2011), the DFE of the standing genetic variation (Orr and Betancourt, 2001; Zhang et al., 2004), and the DFE of variants fixed during adaptation (Orr, 2010). We show that migration-selection balance may be another source of *s*-*h* correlation: especially when RI is low and gene flow rather strong, recessive alleles that are divergently maintained at equilibrium tend to have higher selection coefficients than dominant alleles, even when no such correlation exists *a priori* (fig. 7).

### Linkage

We developed a heuristic approximation for the gff when loci are linked across the genome using two-locus theory (Appendix B), and we showed that it accurately predicts equilibrium differentiation and swamping thresholds when linkage is not too tight (roughly *r/s* ≥ 4). Comparison with individual-based simulations for the human and *Drosophila* genetic maps, shows that our approximations are reasonably accurate for realistic genetic maps, at least when divergently selected loci are not tightly clustered in the genome. For the *Drosophila* genetic map however, where tight linkage is common, we predict swamping at lower migration rates than observed in individual-based simulations, whereas for the human genetic map we can accurately predict swamping thresholds (fig. 9).

### Inference of barriers to gene flow

Our results bear relevance to the inference of barriers to gene flow from genomic data. Recent approaches to quantify gene flow in pairs of diverging populations based on observed genetic variation have accounted for heterogeneity in barrier effects by assuming a demographic model in which the migration rate *m* (interpreted as *m*_*e*_) varies along the genome (Fräısse et al., 2021; Laetsch et al., 2023). However, none of these approaches explicitly models the underlying genetic architecture of divergent selection that is assumed to cause variation in *m*_*e*_ across the genome.

So far, the only approach which has attempted this is the one by Aeschbacher et al. (2017), which combines information about local recombination rates with deterministic population genetic theory to obtain predictions of *m*_*e*_ across the genome. They assume a homogeneous genetic architecture with no dominance, where selected loci occur with a constant density across the genome, and infer the selection density per unit of map length (together with the migration rate) through its effect on observable neutral differentiation. We wonder, naturally, whether our approximations could be fruitfully employed in a similar inferential approach, allowing for drift and heterogeneous barrier architectures while avoiding the assumption that introgressing alleles are rare. Such an approach that explicitly connects variation in *m*_*e*_ across the genome to the numbers and fitness effects of selected loci could allow for more detailed inferences about the genetic architecture of local adaptation and RI, and provide insights about the limits to such inference.

### Limitations of the model

Several important limitations of the model, besides the issues arising from tight linkage, should be noted. Firstly, we have assumed that local fitness is determined by an additive trait under directional selection, considering a history where a rapid polygenic selection response has driven up allele frequencies to near fixation. Alternatively, populations may be under stabilizing selection towards different optima. With stabilizing selection and abundant standing variation, a polygenic selection response may initially only involve subtle changes in allele frequencies (Sella and Barton, 2019; Hayward and Sella, 2022), and there may be considerable genetic redundancy (Yeaman, 2015; Barghi et al., 2020), leading to a scenario that is quite different from the one assumed in this paper. The extent of reproductive isolation that can be maintained when these aspects of polygenic adaptation become important remains unclear and likely requires different approaches.

Secondly, our focus on the maintenance of polygenic local adaptation and the reproductive isolation it causes provides only part of the picture, as we have ignored both the initial polygenic response, and the later build-up of divergence in the face of gene flow. We have considered how a *given* genetic architecture underlying divergent selection results in observable patterns of adaptive differentiation at equilibrium, but remain largely ignorant about what would be a plausible genetic architecture of local adaptation. We have also completely ignored epistasis, which may be an important additional source of selection against hybrids and their descendants (e.g. Dobzhansky-Muller incompatibilities). All these are important topics deserving further study if we are to understand how populations can remain locally adapted when subjected to maladaptive gene flow, and, ultimately, the adaptive processes that could drive the origin of new species.

## Author contributions statement

A.Z, H.S. and C.F. conceived the project, A.Z. conducted mathematical analyses and simulations, A.Z. wrote the manuscript with input and feedback from H.S. and C.F.

## Data availability statement

The authors affirm that all data necessary for confirming the conclusions of the article are present within the article, figures, and supplementary material. Software implementing the numerical methods and individual-based simulations is available at https://github.com/arzwa/MultilocusIsland.

## Acknowledgments

This work was funded by the European Union (ERC BryoFit 101041201 granted to CF). Views and opinions expressed are however those of the author(s) only and do not necessarily reflect those of the European Union or the European Research Council. Neither the European Union nor the granting authority can be held responsible for them. We thank Nick Barton, Nicolas Bierne and four anonymous reviewers for helpful comments on an earlier version of the manuscript that substantially improved the presentation of our work.

## A. Detailed derivation of eq. (4)

Under the assumptions stated above eq. (4), each of the *W* ^(*n*)^ is determined by the allele frequencies and heterozygosities at the selected loci in the mainland and the island populations at the assumed equilibrium. This allows us to determine 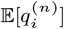, the expected frequency of the locally deleterious allele (in the island) at locus *i* among *n*th generation descendants from a migrant, in terms of the allele frequencies in the mainland and island populations. Indeed, we have the recursive relation 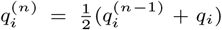, i.e. the expected frequency of the deleterious allele at locus *i* in an *n*th generation descendant is the mean of the corresponding frequencies in the resident population and the (*n* − 1) generation backcrosses. Hence, we have 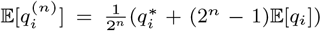. Denoting the selection coefficient at locus *i* for the haploid phase by *s*_*i*1_, the expected relative fitness of an *n*th generation haploid descendant is

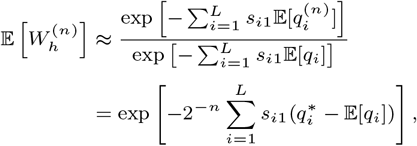

where we have assumed that per-locus selection is sufficiently weak that *O*(*s*^2^) terms can be ignored. For the diploid phase, a similar argument shows that for the (*n* + 1)th generation,

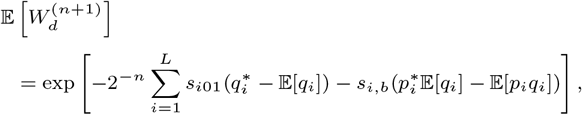

where *s*_*i*01_ and *s*_*i*11_ are the selection coefficients against heterozygotes and homozygotes at locus *i* respectively, and where, analogous to the single-locus model, *s*_*i,b*_ = *s*_*i*11_ − 2*s*_*i*01_. Putting everything together, the approximate gff becomes

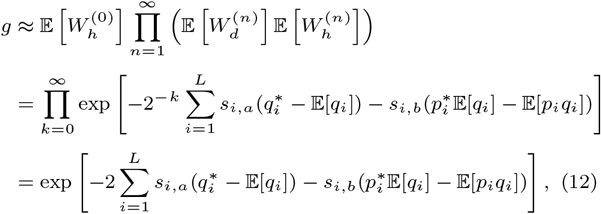

Two remarks are due. Firstly, the gff as derived above yields the effective migration rate at an unlinked neutral locus. We can calculate the gff at a selected locus by making the assumption that it is the same as that of a hypothetical neutral locus at the same location – an assumption which is only expected to work well if selection at the focal locus is sufficiently weak. Hence, if we wish to calculate the effective migration rate for a selected locus in the barrier, say locus *j*, the relevant gff is obtained by excluding index *j* from the sum in eq. (12).

Secondly, we have assumed that migration occurs at the start of the haploid phase, e.g. due to spore dispersal. While the details of when migration occurs in the life cycle do not matter for the single-locus model as long as selection and migration are sufficiently weak (so that the continuous-time limit is appropriate), these details *do* matter for the effective migration rate. This is because, although selection *per locus* is weak (*s* being small), selection against migrant genotypes can be strong (*Ls* being appreciable). Thus, when migration is due to dispersal of gametes (e.g. pollen dispersal), the first generation experiencing selection on the island will be the diploid F1 generation, so that the appropriate gff under the same approximation is 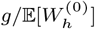. Secondly, when migration occurs at the beginning of the diploid phase (e.g. seed dispersal), the first generation experiencing selection will consist of diploid migrant individuals, so that 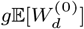 is the appropriate gff, where

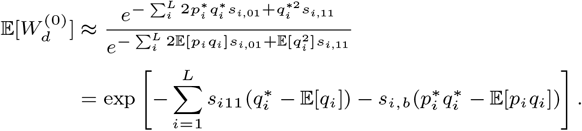

If the haploid, diploid and gametic migration rates are *m*_1_, *m*_2_ and *m*_3_ respectively, the effective migration rate will be (*m*_1_ + 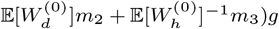.

## B. Accounting for (weak) linkage

To heuristically account for weak linkage between *L* loci, we shall extrapolate from a two-locus model at quasi-linkage equilibrium (QLE). We shall hence first consider the deterministic dynamics of a two-locus haplodiplontic model with mainland-island migration in continuous-time.

We assume a locus *A* with alleles *A*_0_ and *A*_1_ and a linked locus *B* with alleles *B*_0_ and *B*_1_, with recombination between the two loci occurring at rate *r*. We assume arbitrary dominance and no epistasis, with relative Malthusian fitnesses of all possible two-locus genotypes in the two phases given by the following tables

Haploid phase

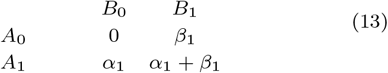

Diploid phase

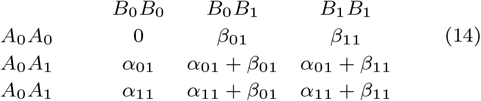

Similar to our notation for the single-locus case, we let *α*_*a*_ = *α*_1_ + *α*_01_ and *α*_*b*_ = *α*_11_ − 2*α*_01_ (and similarly for *β*_*a*_ and *β*_*b*_). Let *x*_*ij*_ and *y*_*ij*_ denote the frequency of the *A*_*i*_*B*_*j*_ haplotype on the island and mainland respectively. The two-locus dynamics in continuous-time are given by

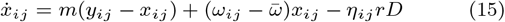

where *D* = *x*_00_*x*_11_ − *x*_01_*x*_10_ is the usual measure of two-locus LD, *ω*_*ij*_ the marginal fitness of the *A*_*i*_*B*_*j*_ haplotype, 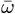 the mean Malthusian fitness and *η*_*ij*_ = 1 when *i* = *j* and −1 otherwise. We write *p*_*A*_ = 1 − *q*_*A*_ = *x*_00_ + *x*_01_ for the frequency of the *A*_0_ allele, and *p*_*B*_ = 1 − *q*_*B*_ = *x*_00_ + *x*_10_ for the frequency of the *B*_0_ allele.

Defining

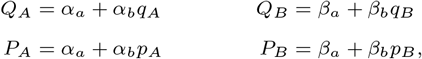

one can find that 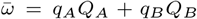 and write the marginal fitnesses as

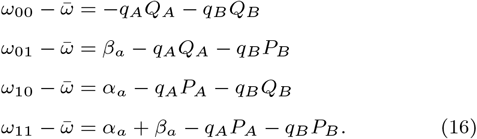

Using eq. (16) with eq. (15), and assuming the mainland is at HWLE with allele frequencies 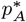 and 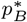 at the two loci, one can derive the dynamics for *p*_*A*_, *p*_*B*_ and *D*:

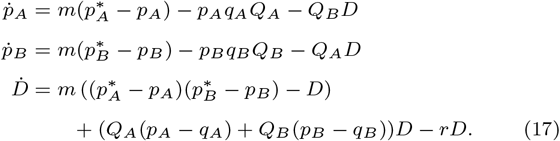

We can use eq. (17) to derive the effective migration rate at a neutral locus linked to a barrier locus maintained at migration-selection balance. Let *A* be the selected locus, and *B* the linked neutral locus. The dynamics of the system are given by eq. (17), where *Q*_*B*_ = 0. Assuming *r* is sufficiently large relative to the strength of selection (linkage is sufficiently weak) so that *D* equilibrates much faster than the allele frequencies (QLE assumption), we can solve the system for *D* at equilibrium to find

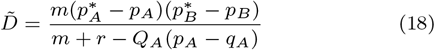

We can plug this into the ODE for the neutral locus to find

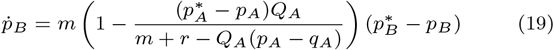

Suggesting that the effective migration rate under the stated assumptions should be

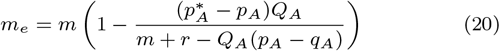

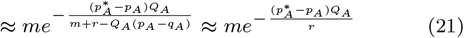

where the second approximation holds well for weak linkage (*r* ≫ *s* ∼ *m*), which we assumed when we derived eq. (19). If we now consider a multilocus system with *L* loci, and assume that the effects of the *L* − 1 other loci on gene flow at a focal locus act multiplicatively, we can approximate the gff at locus *i* as in eq. (5).

