## Supplemental Information for "The genetic architecture of polygenic local adaptation and its role in shaping barriers to gene flow"

### S1. Supplementary figures

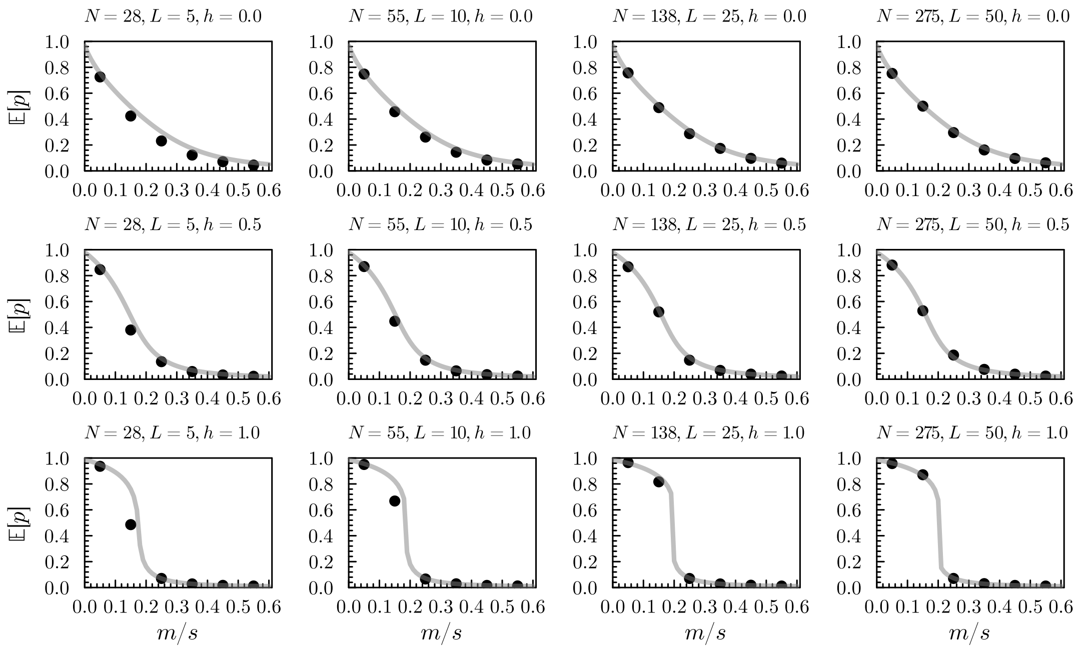


Fig. S1: Comparison of the multilocus diffusion approximation (gray line) against individual-based simulations (black dots). $Ls = 1$ and $N_e s = 5$ for all plots, while $L$ is varied across columns ($L \in [5, 10, 25, 50]$) and $h$ varies over rows ($h \in [0, 0.5, 1]$). We assumed $k = 5$ diploids per haploid individual and set $N = N_e/2k + N_e$ so that the desired $N_e = N_e s/(Ls/L)$ is obtained. Allele frequencies for the individual-based simulations are obtained by simulating for 110000 generations, sampling every 5 generations after discarding the first 60000, and averaging across loci. For each $L$ we simulate $n$ replicates so that $nL = 50$. The mutation rate was set to $u = 0.005s$.

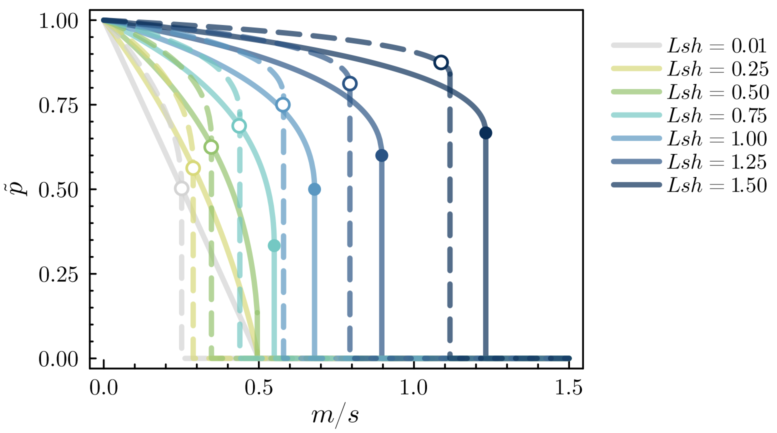


Fig. S2: Equilibrium differentiation and swamping thresholds for the deterministic multilocus model, comparing different degrees of dominance on the basis of $Lsh$. The dashed line shows results for $h = 1$ (recessive local adaptation), whereas the solid line shows results for $h = 0.5$.

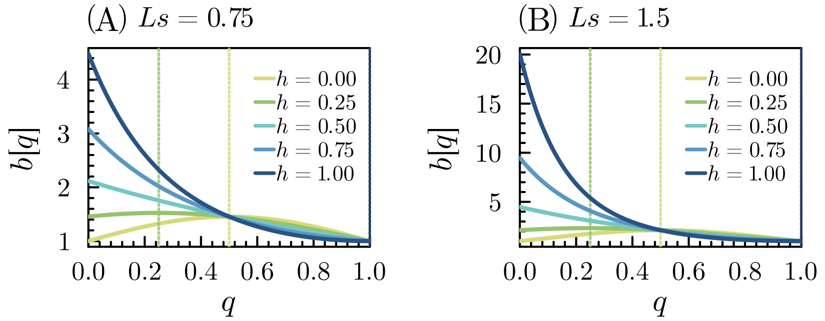


Fig. S3: Barrier strength ($b = g^{-1}$) as a function of the deleterious allele frequency on the island (i.e. differentiation) for different degrees of dominance. Deterministic predictions are shown based on eq. (6) for (A) $Ls = 0.75$ and (B) $Ls = 1.5$. Other paremeters are as in fig. 3

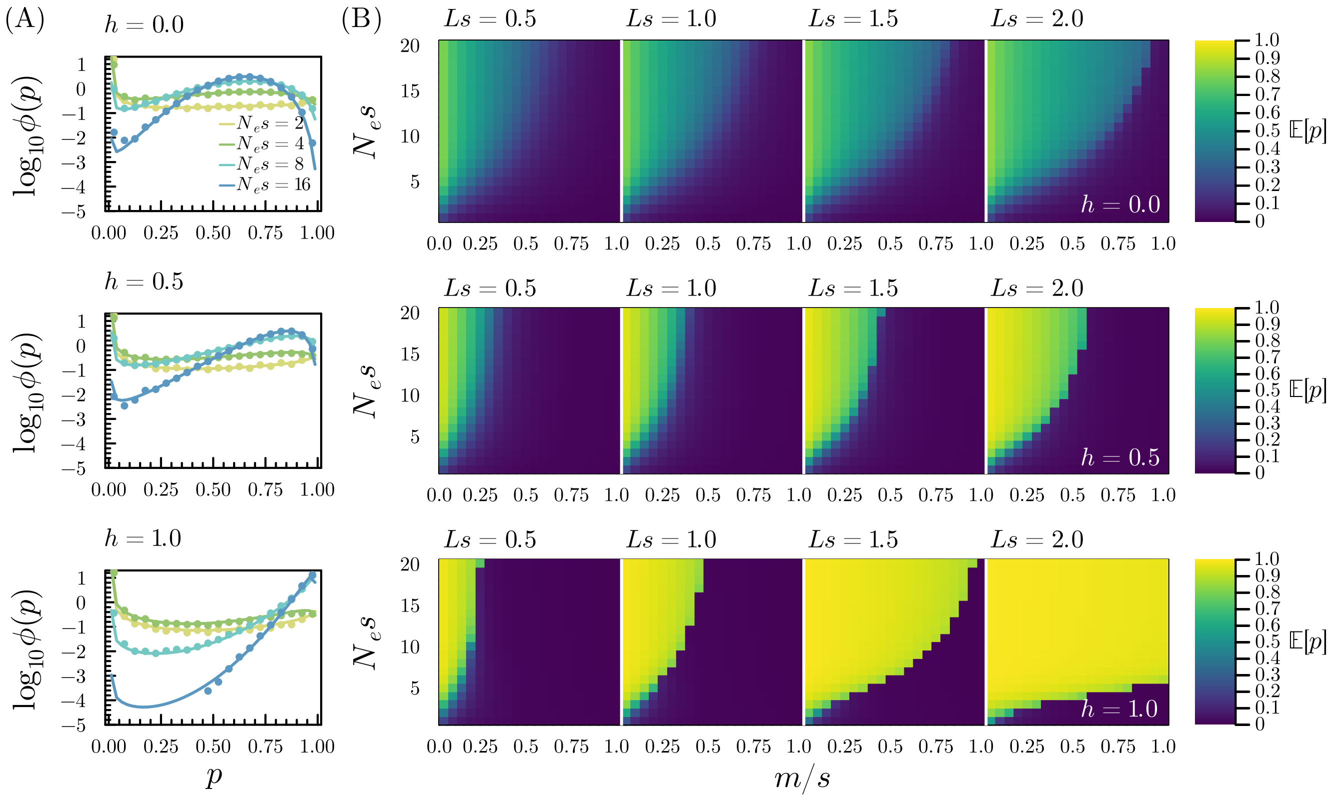


Fig. S4: Genetic drift reduces the strength of a homogeneous multilocus barrier to gene flow. (A) Predicted allele frequency distributions for a single-locus in the diploid multilocus model with homogeneous selective effects for different values of $N_e$ and $h$ (dominance coefficient of the invading alleles). Lines show the numerical approximations based on the diffusion theory, dots show results from individual-based simulations, based on taking a sample every 10 generations for 50000 generations after an initial 10000 generations to reach equilibrium. In these simulations, $s = 0.02$, $Ls = 1$ and $u/s = 0.005$. (B) The expected frequency of the locally beneficial allele ($\mathbb{E}[p]$) is shown as a function of $N_e s$ and $m/s$ for different values of $Ls$ (from left to right, $Ls = 0.5, 1, 1.5, 2$) and $h$ (from top to bottom, $h = 0, 0.5, 1$), computed using the numerical approximation. All results assume $s = 0.02$ and $u/s = 0.005$.

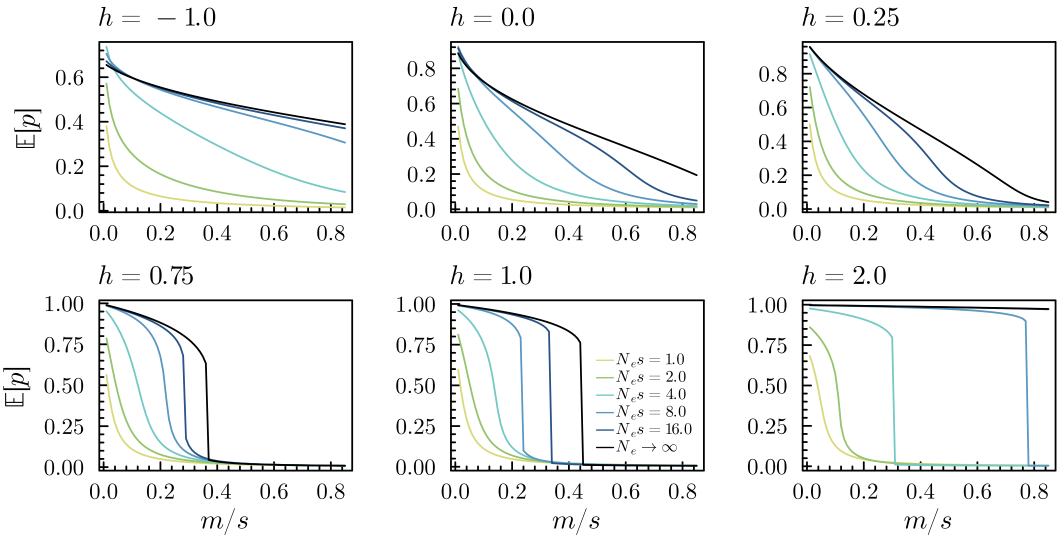


Fig. S5: Effect of drift on equilibrium differentiation and swamping thresholds for a range of dominance values $h$, ranging from overdominant local adaptation $h = -1$ (hybrids have an advantage), to underdominant local adaptation $h = 2$ (hybrids perform worse than mainland individuals on the island). All results use $L = 40, Ls = 0.8, k = 5$.

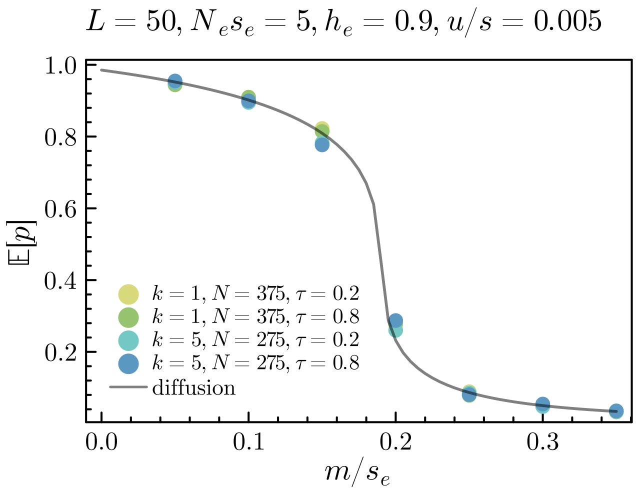


Fig. S6: Effective parameters accurately describe equilibrium dynamics for haplodiplontic populations when selection is sufficiently weak. The line shows the numerical prediction of the locally beneficial allele frequency on the island for increasing strength of migration relative to *effective* selection. The dots show results from individual based simulations with different degrees of haploid vs. diploid selection and different relative sizes of the haploid and diploid population, keeping $N_e, s_e$ and $h_e$ however constant. Simulation results are based on 110000 generations, where we sampled every 10th generation after discarding the first 10000 generations.

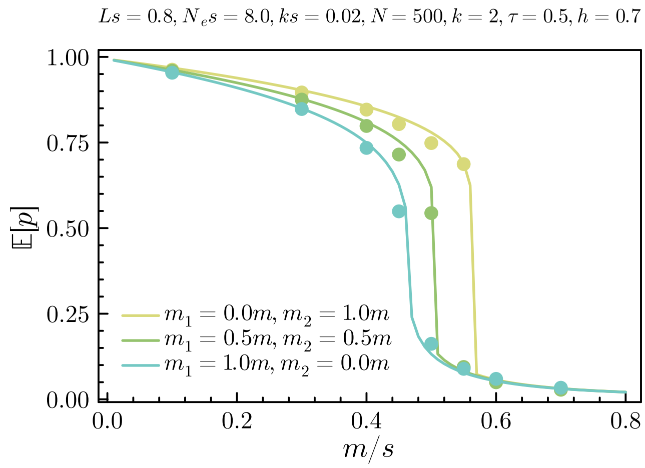


Fig. S7: Migration in the diploid phase of a haplodiplontic life cycle with selection in both phases leads to stronger barriers to gene flow. The lines show predictions from the multilocus diffusion theory, whereas the dots show results from individual-based simulations (taking a sample every fifth generation during 20000 generations after discarding the first 5000 generations). The migration rates in the haploid and diploid stage are $m_1$ and $m_2$ respectively.

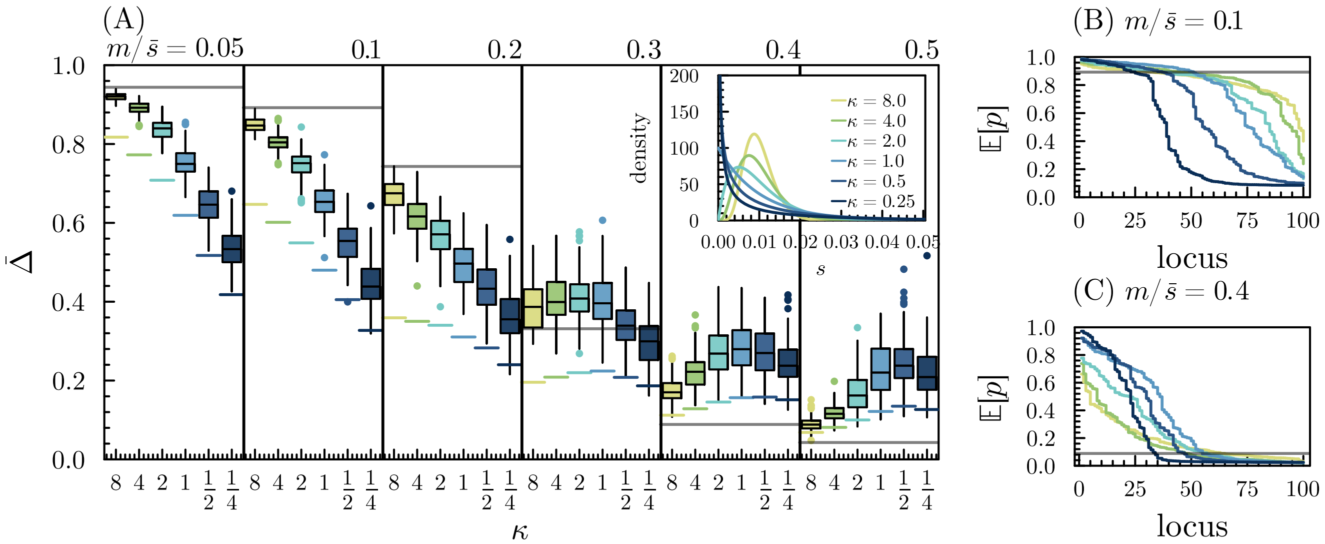


Fig. S8: As in fig. 5, but where we assume $h_i \sim \text{Beta}(1, 1)$ independently for each locus in the barrier (instead of assuming $h_i = 1/2$ for all $h$ as in fig. 5). The colored lines show the average differentiation across the barrier predicted using single locus theory $\sum_i \mathbb{E}[p_i|s_i]/L$, averaged over replicate simulations (i.e. predictions accounting for heterogeneity, but not LD).

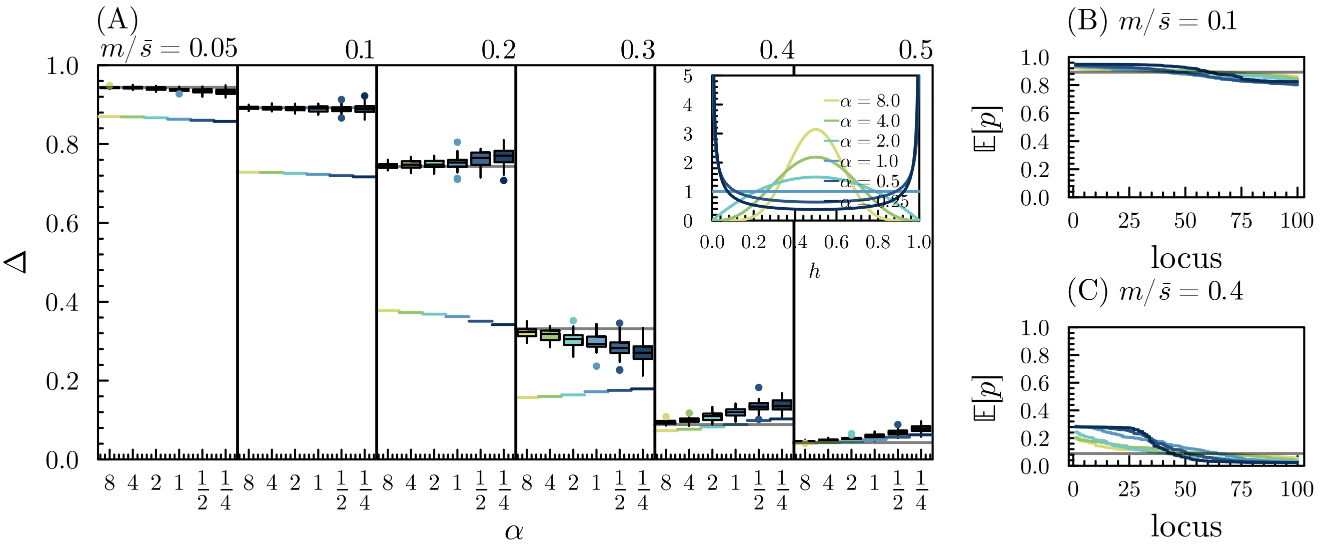


Fig. S9: As in fig. S8, but now keeping the selection coefficient fixed at $\bar{s} = 0.01$ and using randomly sampled dominance coefficients, from a symmetric Beta distribution with parameter $\alpha$. Again, $L = 100, N_e s = 10, u/s = 0.005$.

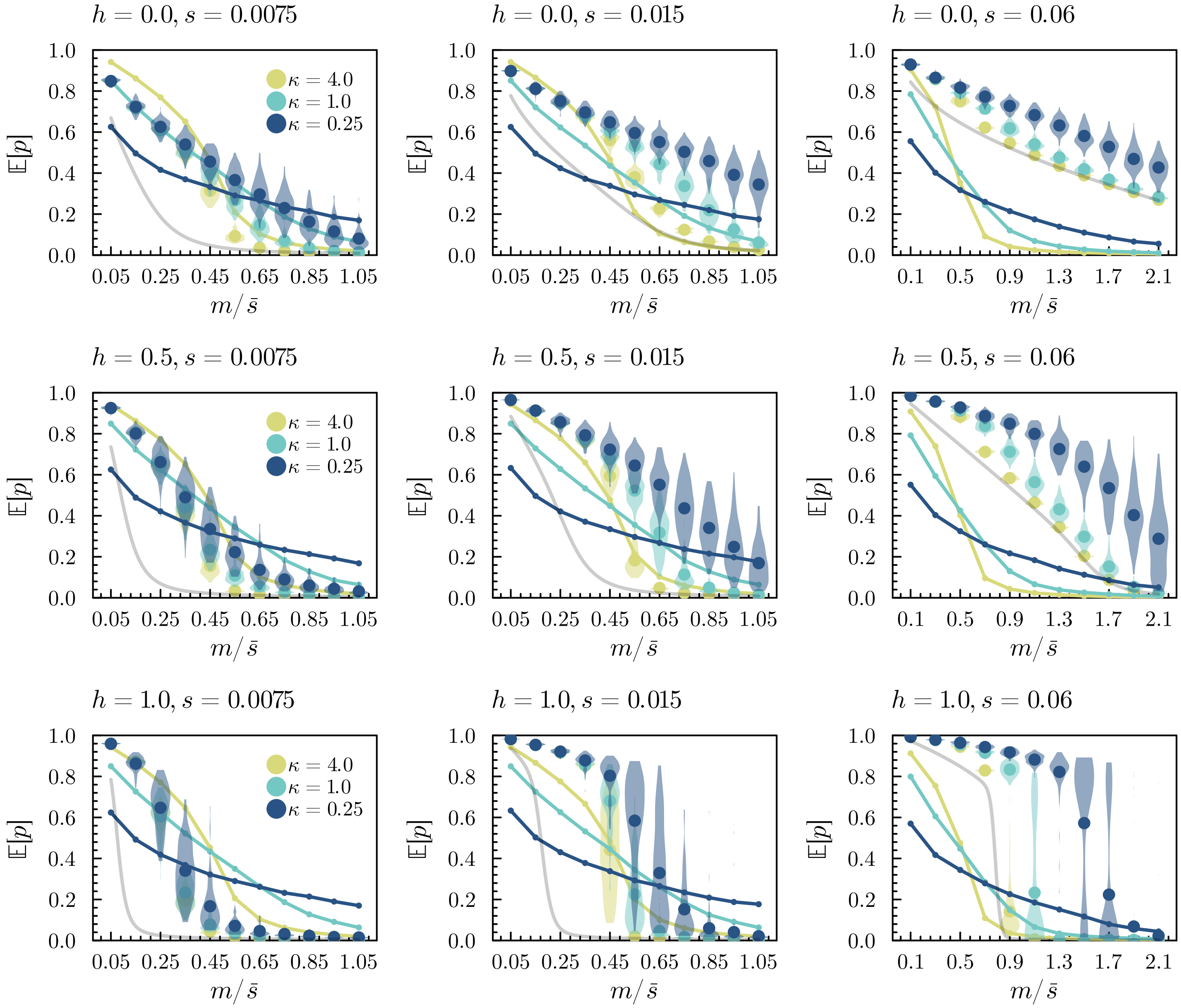


Fig. S10: The effect of barrier heterogeneity on differentiation at a focal locus. The violin plots show the distribution of $\mathbb{E}[p]$ at a focal locus with $s = \bar{s}/2 = 0.0075$ (left column), $s = \bar{s} = 0.015$ (middle column) or $s = 4\bar{s} = 0.06$ (right column) and different assumed dominance coefficients (rows) across 100 replicate simulations of a polygenic barrier with $L\bar{s} = 1.5$ and $L = 100$ in which this locus is embedded, for different values of $m/\bar{s}$ and different values of $\kappa$, where $s_i \sim \text{Gamma}(\kappa, \kappa/\bar{s})$ and $h_i \sim \text{Uniform}(0, 1)$ (recall that $\kappa^{-1} = \text{Var}[s]/\bar{s}^2$). The dots show the average differentiation at the focal locus across the 50 replicates, whereas the lines show $\bar{\Delta}$, i.e. the average expected differentiation across the $L$ loci in the barrier. The gray line shows the associated single-locus prediction for the focal locus. We assume $N_e s = 20$ and $u = s/200$.

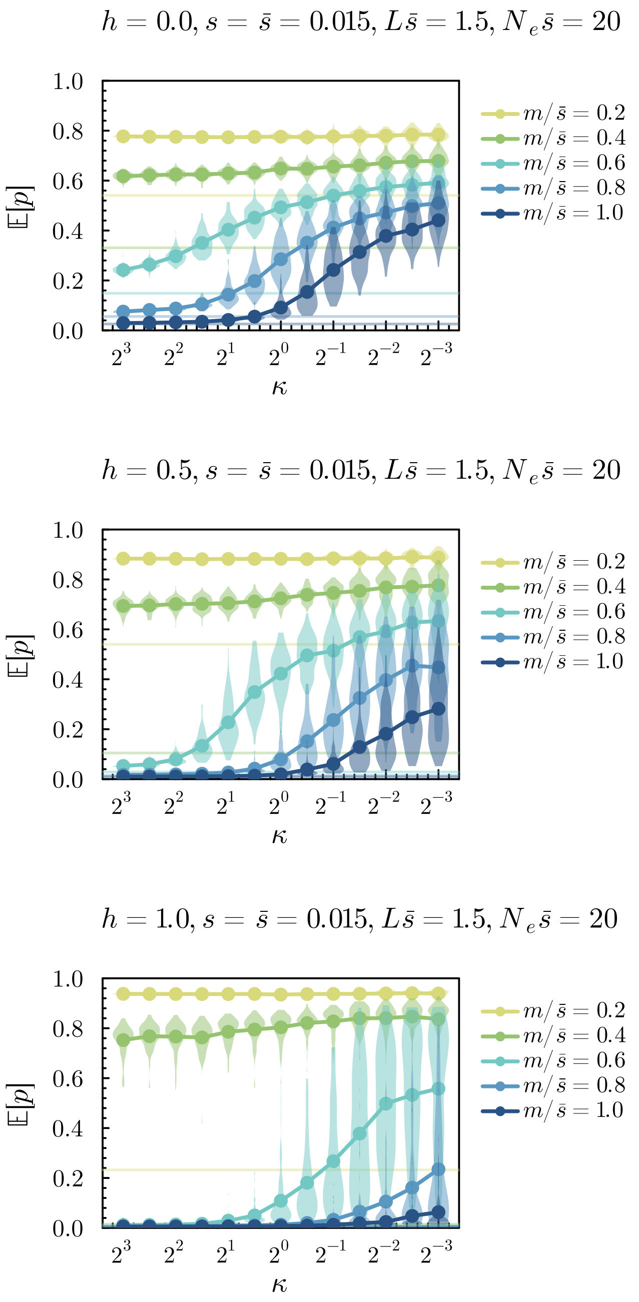


Fig. S11: The effect of barrier heterogeneity on differentiation at a focal locus. The violin plots show the distribution of the expected differentiation at a dominant, additive or recessive locus (from top to bottom) with selection coefficient $s = \bar{s} = 0.015$ embedded in a random heterogeneous barrier with $s_i \sim \text{Gamma}(\kappa, \kappa/\bar{s})$ and $h_i \sim \text{Uniform}(0, 1)$, estimated using 100 replicate simulations (recall that $\kappa^{-1} = \text{Var}[s]/\bar{s}^2$). The dots show the mean expected differentiation across replicates. The horizontal lines show the single-locus predictions for the focal locus at the relevant value of $m/\bar{s}$. Other parameters are as in fig. S10.

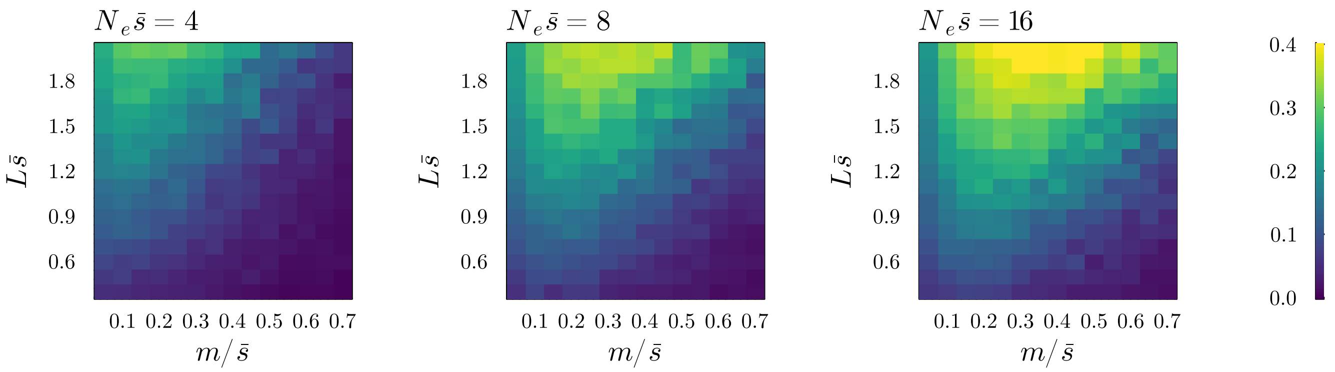


Fig. S12: Average difference in predicted allele frequency for the single locus vs. multilocus model. We assume $L = 100$, $s_i \sim \text{Exponential}(\bar{s})$ and $h_i \sim \text{Beta}(1,1)$ for $i = 1, \dots, L$. We show $\frac{1}{L}\sum_i^L |\mathbb{E}[p_{i,L}] - \mathbb{E}[p_{i,\text{single}}]|$ where $p_{i,L}$ and $p_{i,\text{single}}$ are the equilibrium frequency of the locally beneficial allele at locus $i$ in the multilocus model and single-locus model respectively. The results are averaged across 10 random $L$-locus barriers. We show results for different strengths of genetic drift ($N_e\bar{s}$). Note that values of $L\bar{s}$ range from 0.5 to 2 ($y$-axis).

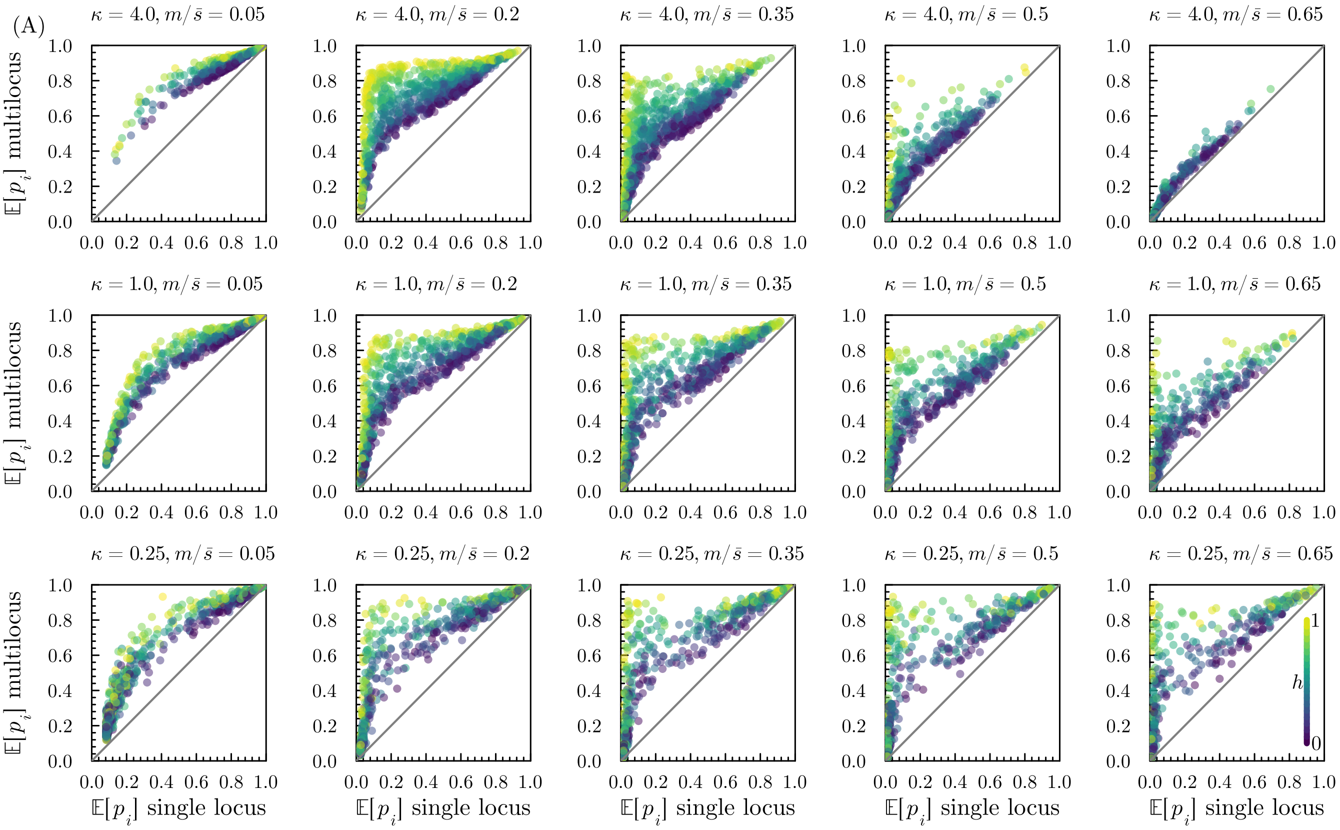


Fig. S13: As in fig. 6, but varying the extent of barrier heterogeneity ($\kappa = 4, 1, 1/4$, rows). Results are shown for $L\bar{s} = 1, \bar{s} = 0.01$. Recall that $\kappa^{-1} = \text{Var}[s]/\bar{s}^2$.

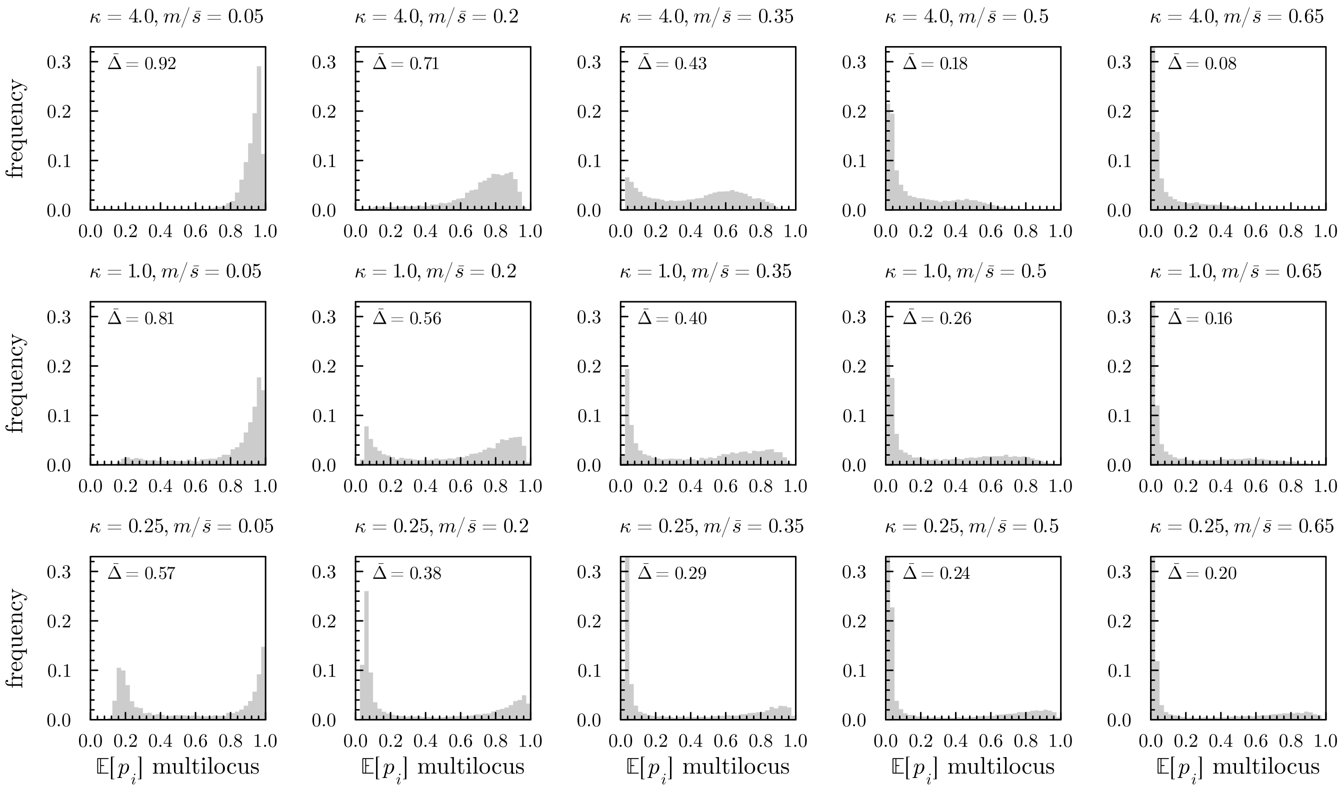


Fig. S14: Distributions of expected equilibrium differentiation at individual barrier loci for the simulations shown in fig. S13.

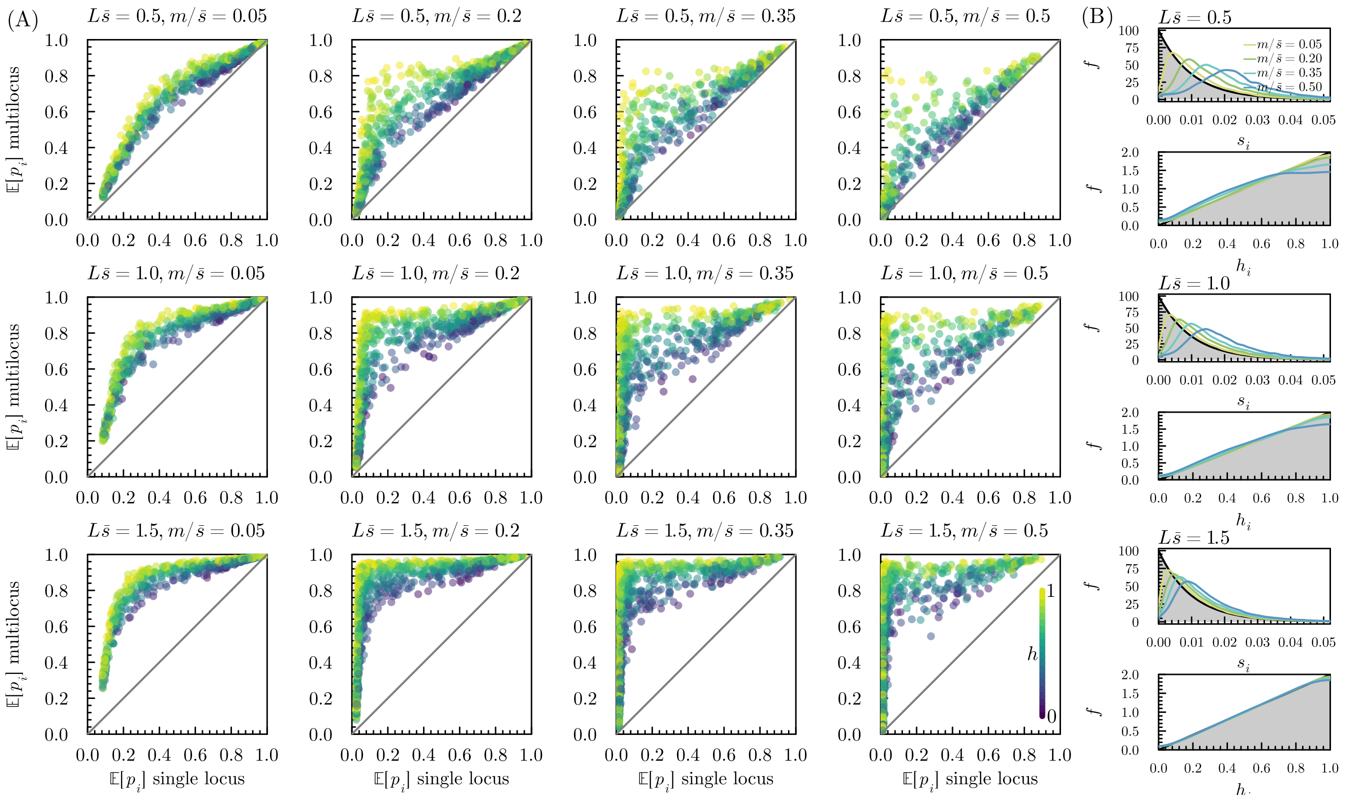


Fig. S15: As in fig. 6, but with $h \sim \text{Beta}(2, 1)$ (so that $\mathbb{E}[h] = 2/3$).

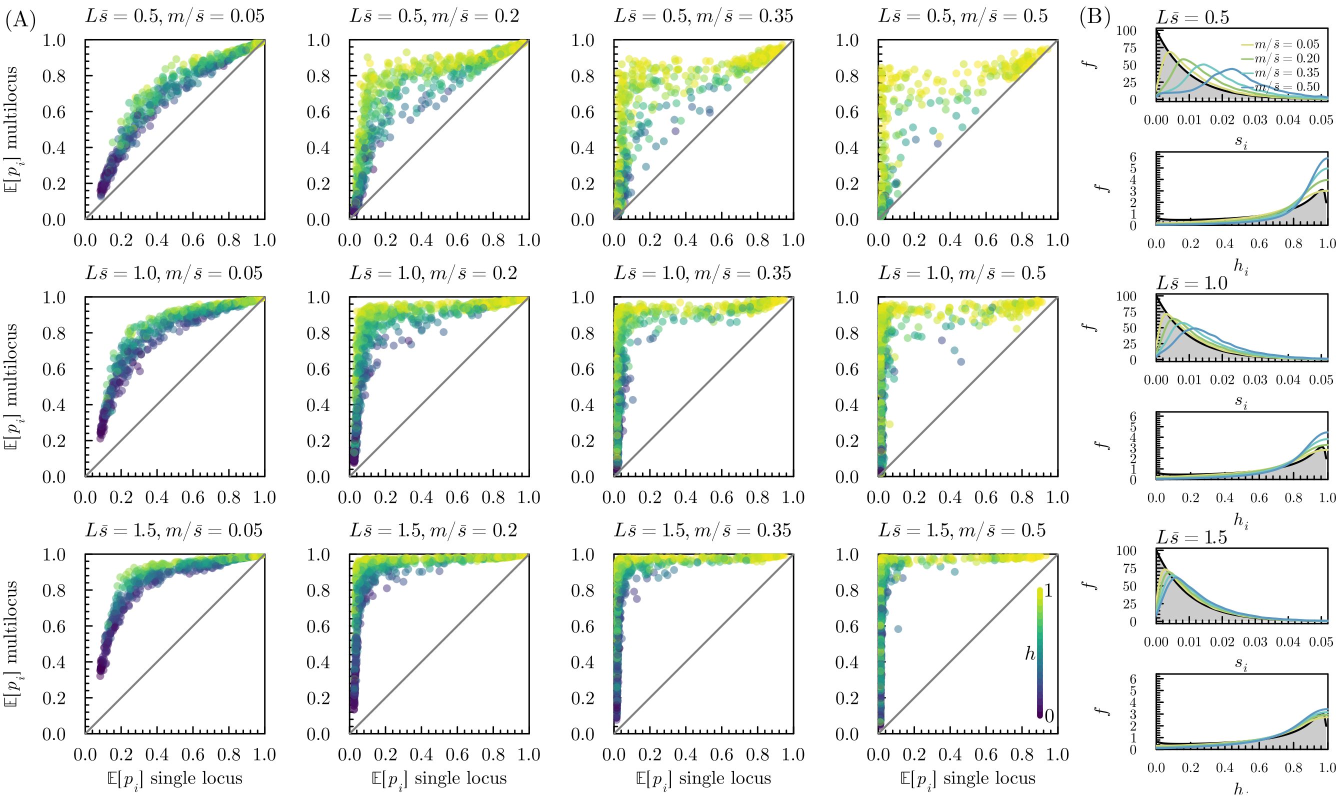


Fig. S16: As in fig. 6, but for the logistic regression model (with $\bar{s} = 1, \kappa = 1, a = 7.2, b = 1.2, \mathbb{E}[h] \approx 2/3, \sigma = 1$; see section S2.6).

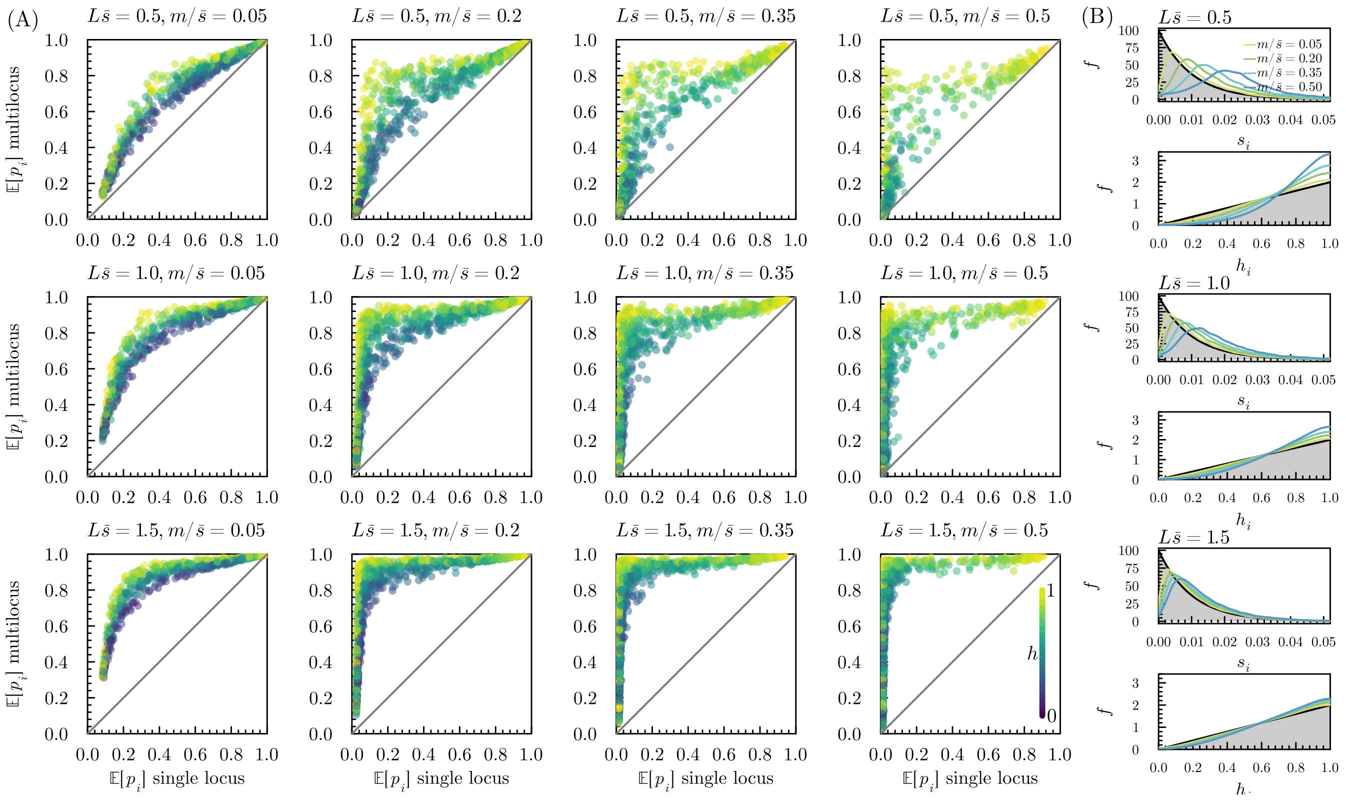


Fig. S17: As in fig. 6, but for the CK94 model (with $\bar{s} = 0.01, \kappa = 1$ and $\mathbb{E}[h] = 2/3$, yielding $K = 50$; see section S2.6).

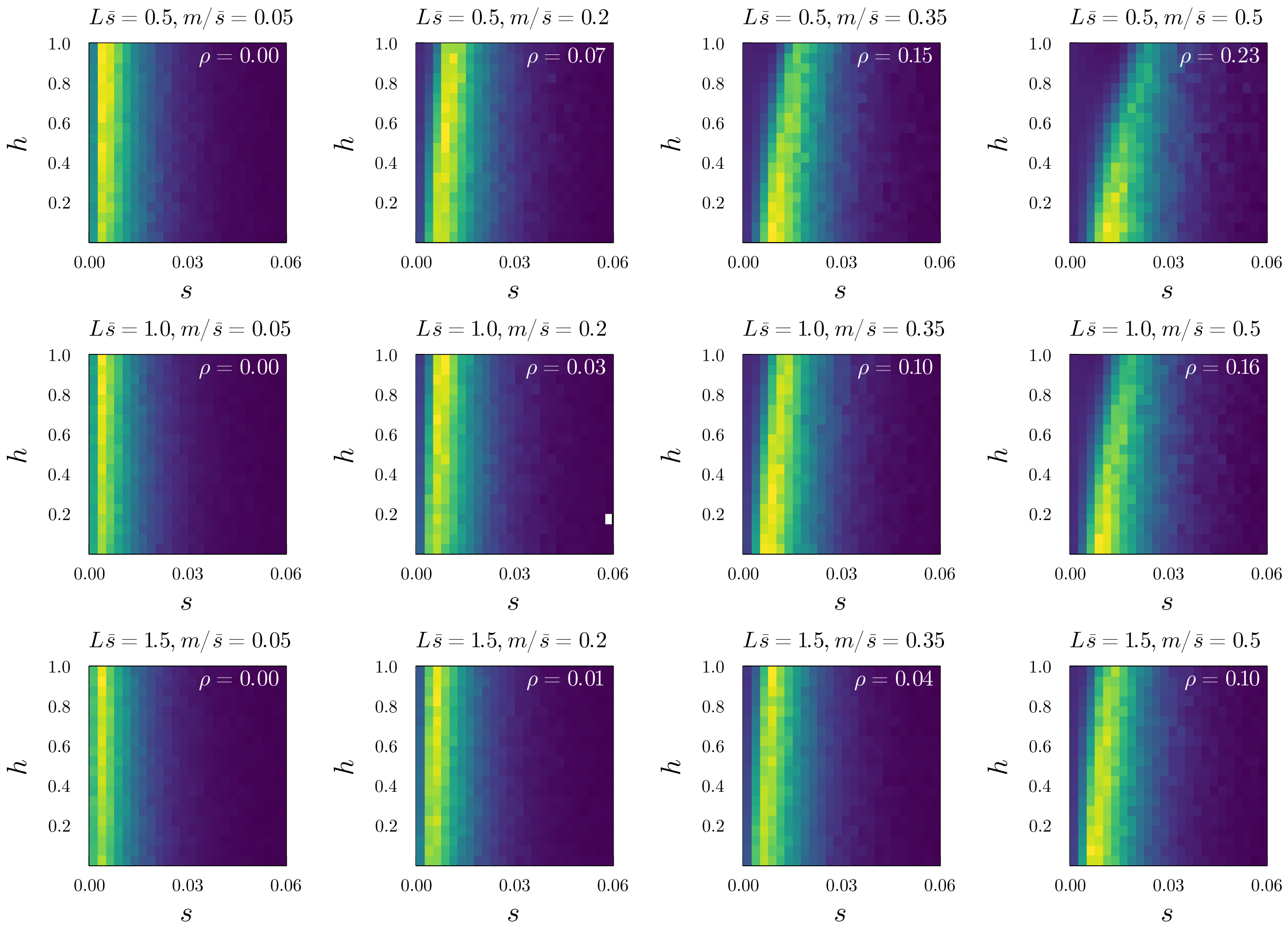


Fig. S18: Monte Carlo approximation to the joint probability distribution of $s$ and $h$ conditional on observing a divergent allele on the island (eq. (8)) for the DFE model with independent selection and dominance coefficients (see fig. 6 and section S2.6). Deep blue designates regions of low probability density, bright yellow regions of high probability density. The estimated correlation $\rho$ between $s$ and $h$ is shown in the upper right corner.

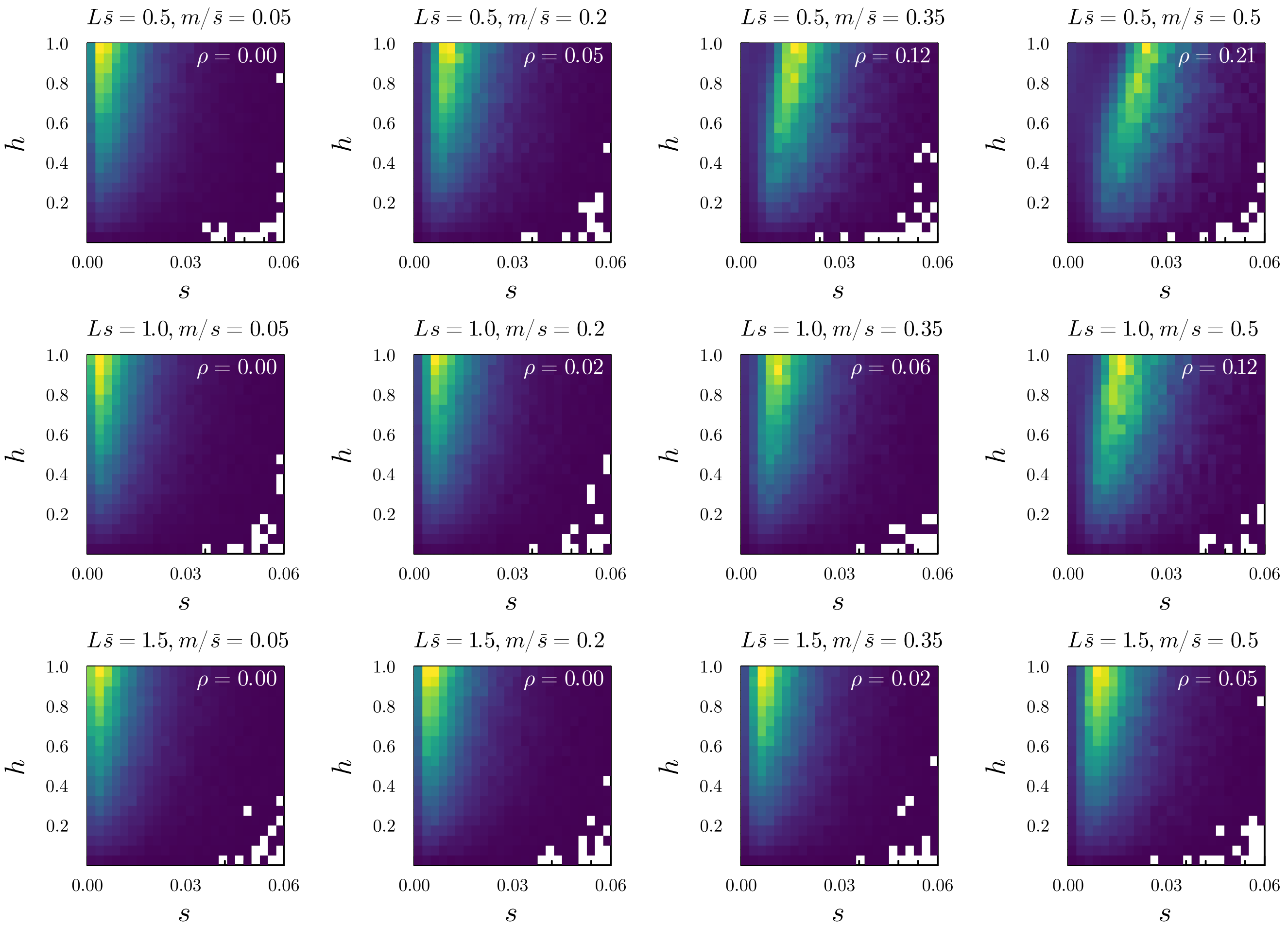


Fig. S19: As in fig. S18, but with $h \sim \text{Beta}(2, 1)$ (see fig. S15 and section S2.6).

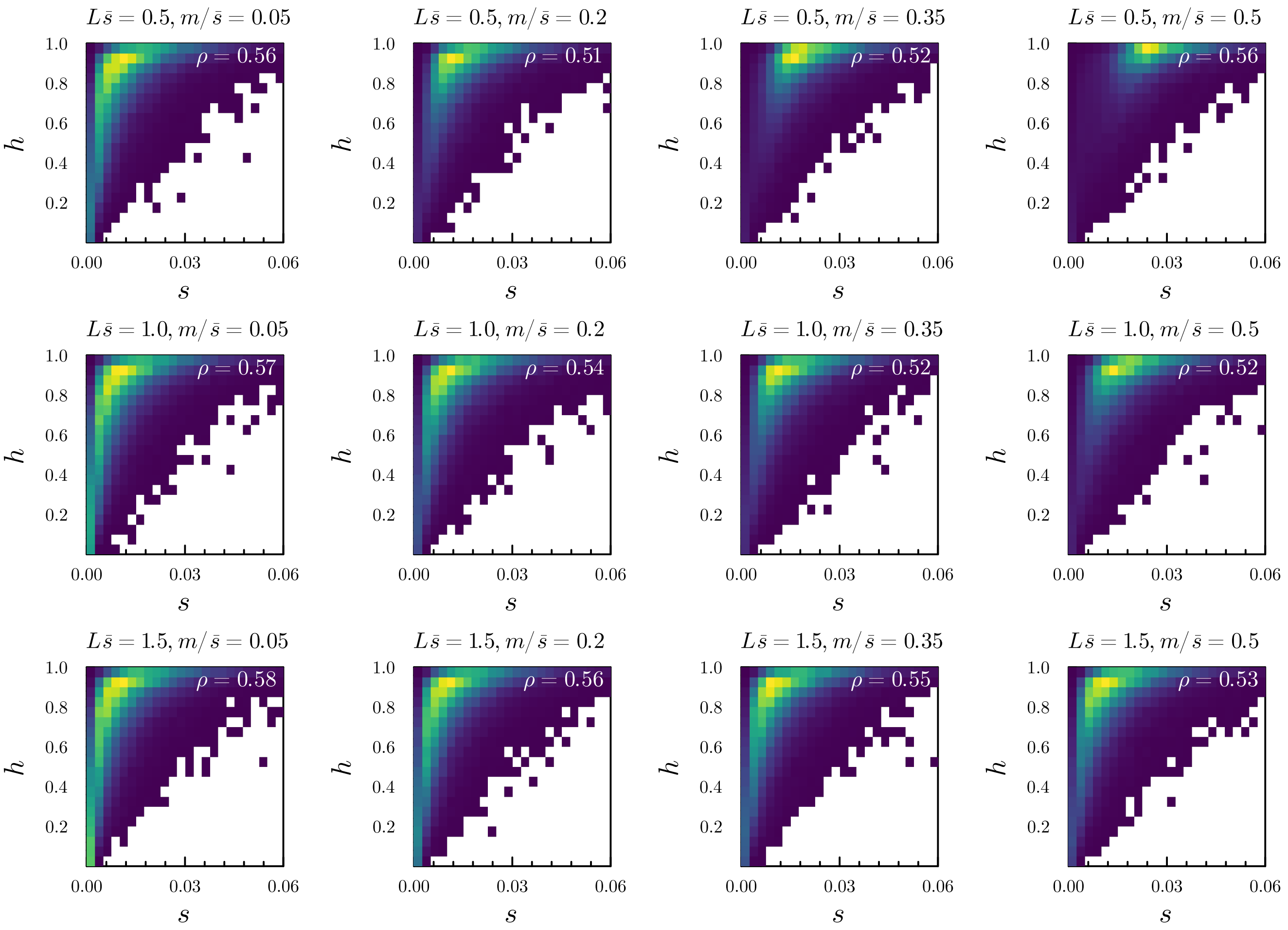


Fig. S20: As in fig. S18, but for the logistic model (see fig. S16 and section S2.6).

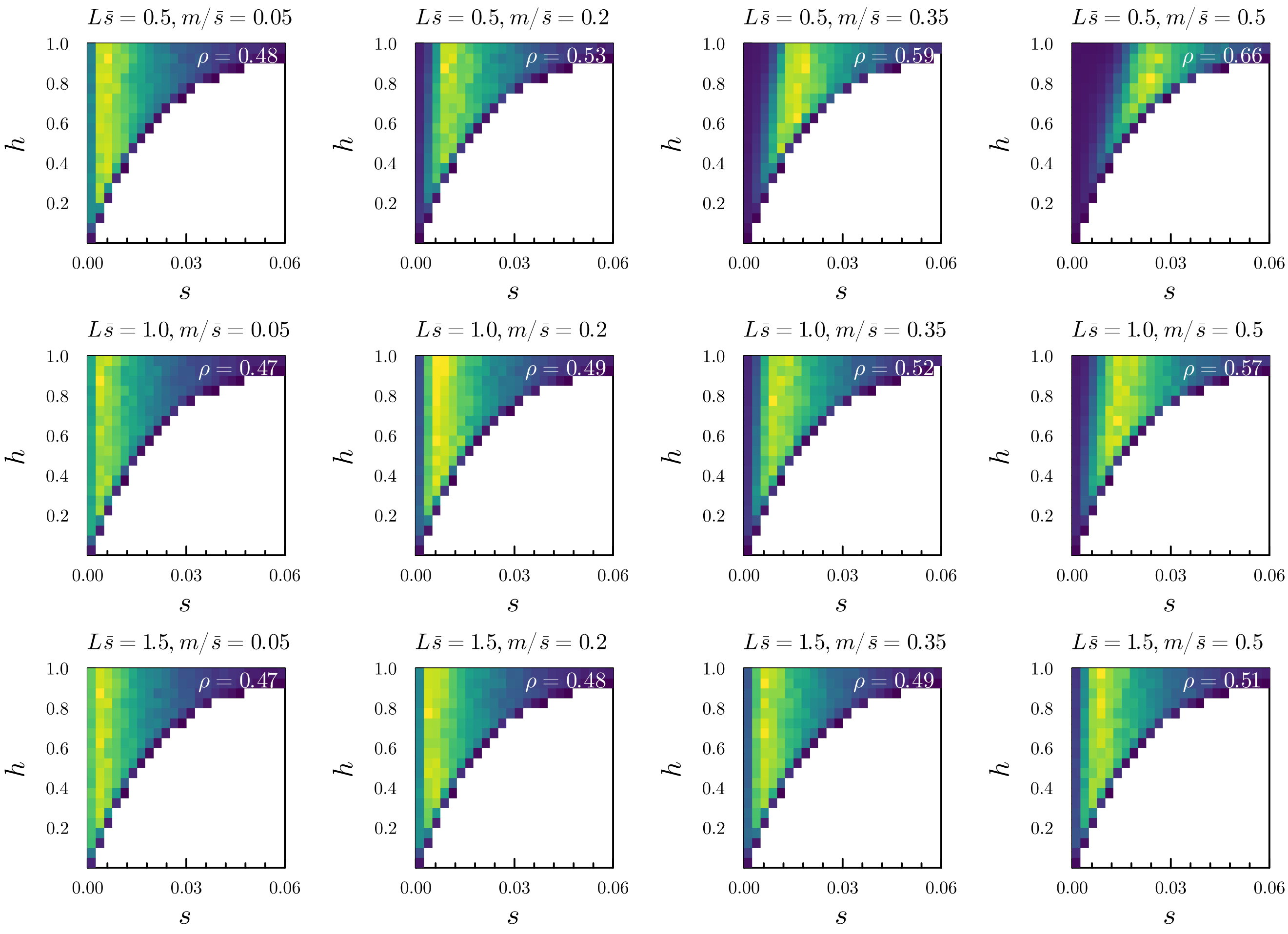


Fig. S21: As in fig. S18, but for the CK94 model (see fig. S17 and section S2.6).

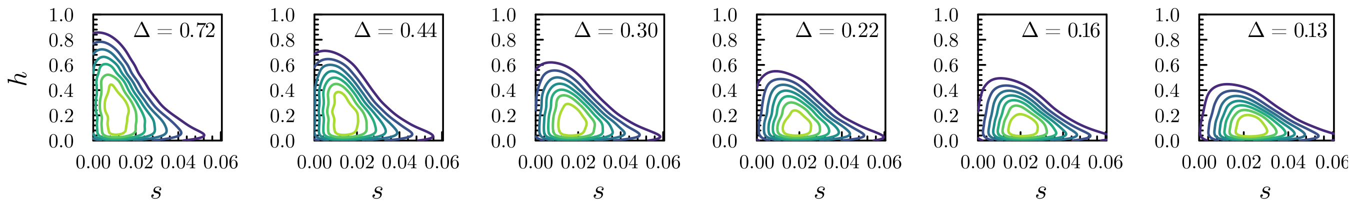


Fig. S22: As in fig. 7, but for the CK94* model with a negative correlation between $s$ and $h$, see section S2.6. $m/s$ values from left to right are $0.05, 0.20, 0.35, 0.50, 0.65$ and $0.80$, as in fig. 7.

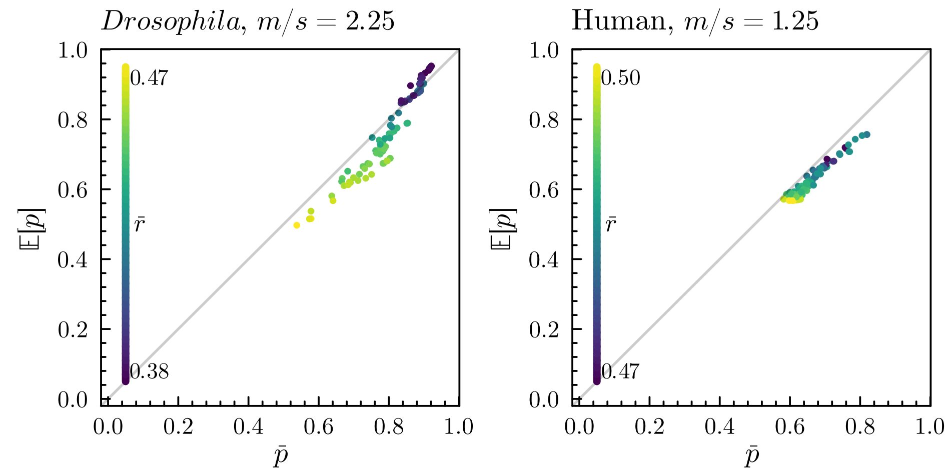


Fig. S23: Scatter plots as in the left panel of figure fig. 9, for a single value of $m/s$ for both the *Drosophila* and Human genetic maps. Dots are colored by the average pairwise recombination rate for the associated locus ($\bar{r_i} = \sum_{j=1}^{L} r_{ij}/L$). Note that we assume equal effect loci, so that $\bar{r} \propto \bar{r}/s$ (which is the key parameter determining the barrier strength).

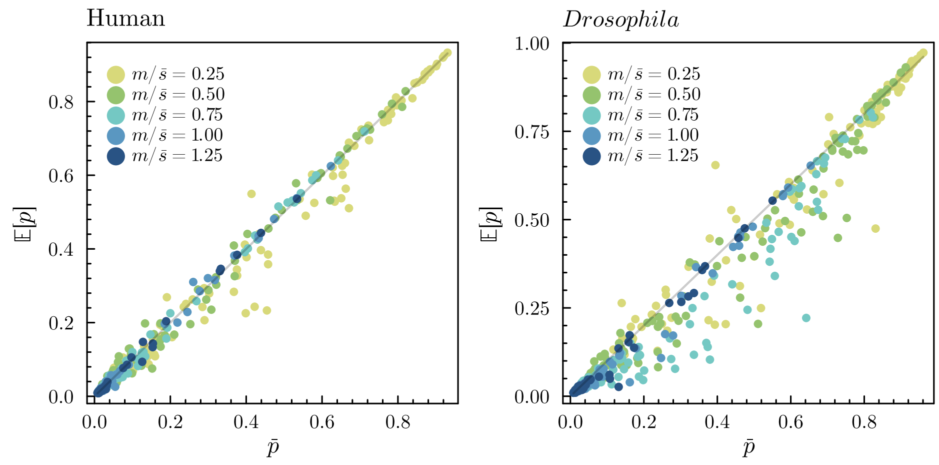


Fig. S24: Comparison of predicted allele frequencies ($\mathbb{E}[p]$) against observed average allele frequencies in individual-based simulations ($\bar{p}$) for a heterogeneous $L$-locus barrier, where the loci are randomly scattered across the human (left) and *Drosophila* (right) genetic maps, for five different migration rates ($m/\bar{s}$ values). Each dot corresponds to a locus. We assume $L\bar{s} = 1.0$, $L = 100$, $N_e\bar{s} = 10$, $s_i \sim \text{Exponential}(\bar{s})$, $h_i \sim \text{Uniform}(0, 1)$, $u/\bar{s} = 1/100$.

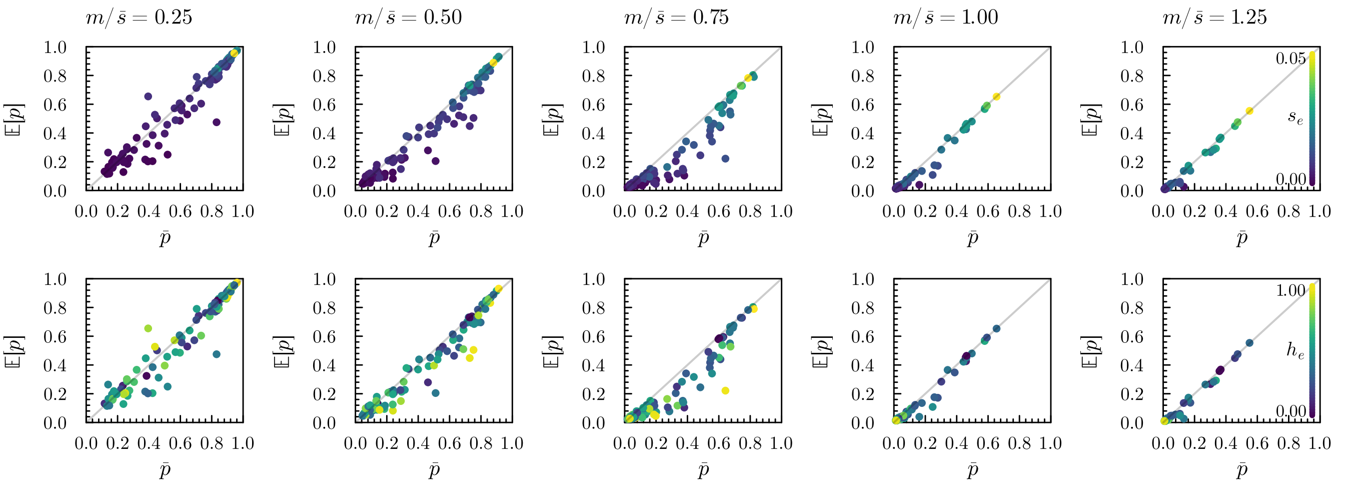


Fig. S25: As in fig. S24, focusing on the *Drosophila* case and plotting the results for every $m/\bar{s}$ value separately, while coloring the dots by effective selection $s_e$ (top row) and effective dominance $h_e$ coefficient (bottom row).

### S2. Supplementary Information

#### S2.1. Single locus allele frequency dynamics for weak selection

Consider a single locus in a population of organisms with a haplodiplontic life cycle. We assume a finite number $n$ of alleles exist at the locus. Let $p_i$, $i \in [0..n-1]$, denote the frequency of the $i$th allele ($A_i$). Ignoring mutation and migration for now, the change in allele frequency of an allele $A_i$ throughout the life cycle is assumed to take the form:

$$\underbrace{p_i \underset{\text{haploid selection}}{\longrightarrow} p_i^* \underset{\text{gametogenesis}}{\longrightarrow} p_i^*}_{\text{haploid (gametophytic) phase}} \underset{\text{syngamy}}{\longrightarrow} \underbrace{p_i^* \underset{\text{diploid selection}}{\longrightarrow} p_i' \underset{\text{meiosis}}{\longrightarrow} p_i'}_{\text{diploid (sporophytic) phase}}, \tag{1}$$

where we have assumed that gametogenesis, syngamy and spore formation do not affect the allele frequencies[1]. Generally, migration could take place at any stage in the life cycle, for instance right after meiosis (e.g. dispersal of meiospores in bryophytes and Fungi), at the end of the haploid phase (e.g. gamete dispersal in algae), early in the diploid phase (e.g. seed dispersal in spermatophytes) or at the level of adult diploids (e.g. migration in animals).

Let the relative haploid fitness of a haploid individual carrying allele $i$ be $1 + \epsilon s_i$, defined as the relative contribution to the diploid (sporophytic) generation within the haploid (gametophytic) generation. Similarly, we let $1 + \epsilon s_{ij}$ denote the relative fitness of a diploid individual with genotype $A_i A_j$. The allele frequency change over a single generation is determined by the following dynamical system:

$$\begin{aligned} p_i^* &= \frac{w_{h,i}}{\bar{w}_h(p)} p_i \\ p_i' &= \frac{w_{d,i}(p^*)}{\bar{w}_d(p^*)} p_i^* \qquad 1 \le i \le n, \end{aligned} \tag{2}$$

where $p = (p_1, p_2, \dots, p_n)$ (and similarly for $p^*$). The marginal fitnesses $w_{h,i}$ and $w_{d,i}$ associated with allele $A_i$ in the haploid (gametophytic) and diploid (sporophytic) phase respectively are

$$\begin{aligned} w_{h,i} &= 1 + \epsilon s_i \\ w_{d,i}(p) &= \sum_j (1 + \epsilon s_{ij}) p_j = 1 + \epsilon \sum_j s_{ij} p_j := 1 + \epsilon \bar{s}_{d,i}. \end{aligned}$$

The mean fitnesses in the gametophytic and sporophytic phases are

$$\begin{aligned} \bar{w}_h(p) &= \sum_i (1 + \epsilon s_i) p_i := 1 + \epsilon \bar{s}_h \\ \bar{w}_d(p) &= \sum_i \sum_j (1 + \epsilon s_{ij}) p_i p_j := 1 + \epsilon \bar{s_d} \end{aligned}$$

The allele frequency change over a single alternation of generations for the dynamical system defined in eq. (2) has the usual form

$$\Delta p_i = \frac{\bar{w}_i - \bar{w}}{\bar{w}} p_i \tag{3}$$

Where, from eq. (2), we have

$$\bar{w} = \left(1 + \epsilon \sum_j \sum_k \frac{(1+\epsilon s_j) p_j (1 + \epsilon s_k) p_k}{(1 + \epsilon \bar{s}_h)^2} s_{jk}\right)(1 + \epsilon \bar{s}_h) = 1 + \epsilon \bar{s}_h + \frac{\epsilon \bar{s}_s}{(1+\epsilon \bar{s}_h)^2} + O(\epsilon^2) \tag{4}$$

and

$$\bar{w_i} = \left(1 + \epsilon \frac{\sum_j s_{ij}(1 + \epsilon s_j) p_j}{1 + \epsilon \sum_k s_k p_k}\right)(1 + \epsilon s_i) = 1 + \epsilon s_i + \frac{\epsilon \bar{s}_{s,i}}{(1 + \epsilon \bar{s}_h)} + O(\epsilon^2),$$

so that

$$\bar{w_i} - \bar{w} = \epsilon(s_i - \bar{s}_h) + \frac{\epsilon}{(1+\epsilon \bar{s}_h)^2}(\bar{s}_{d,i} - \bar{s}_d) + O(\epsilon^2). \tag{5}$$

Assuming the intensity of selection per generation is weak (all $s$ are small) and that $\epsilon$ measures the generation time, we obtain a continuous-time model of allele frequency change by considering the per-generation change in allele frequency and taking the limit as $\epsilon$

[1] Note that we use $p_i^*$ to label the allele frequency after haploid selection, whereas in the main text we use this to denote the mainland allele frequency. Here we are not considering migration however.

goes to zero, specifically, plugging eq. (5) and eq. (4) in eq. (3), we get

$$\dot{p_i} = \lim_{\epsilon \to 0} \frac{\Delta p_i}{\epsilon} = \big[(s_i - \bar{s}_h) + (\bar{s}_{d,i} - \bar{s}_d)\big] p_i$$
$$= (\bar{s}_i - \bar{s}) p_i, \tag{6}$$

where

$$\bar{s}_i = s_i + \bar{s}_{d,i} = s_i + \sum_j s_{ij} p_j$$
$$\bar{s} = \bar{s}_h + \bar{s}_d = \sum_i s_i p_i + \sum_i \sum_j s_{ij} p_i p_j. \tag{7}$$

This has the same form as the classical diploid or haploid continuous-time model of allele frequency change[2] but with marginal and mean Malthusian fitnesses given by $\bar{s}_i$ and $\bar{s}$ respectively.

As an example, consider the biallelic case with alleles $A_0$ and $A_1$, so that $s_0 = s_{00} = 0$, $p_0 = p$ and $p_1 = q$. We shall always assume $s_{ij} = s_{ji}$. Importantly, note that in contrast with the main text, we here assume $w_1 = e^{s_1}$, $w_{01} = e^{s_{01}}$, and $w_{11} = e^{s_{11}}$ (and *not* $w_1 = e^{-s_1}, e^{-s_{01}}$ and $e^{-s_{11}}$ as we do above). We have from eq. (7) $\bar{s}_1 = s_1 + s_{01}p + s_{11}q$ and $\bar{s} = s_1 q + 2s_{01}pq + s_{11}q^2$. Some algebra shows that we can write the ODE for the frequency of the selected allele ($A_1$) as

$$\dot{q} = (\bar{s}_1 - \bar{s})q = pq(s_a + s_b q) \tag{8}$$

where $s_a = s_1 + s_{01}$ and $s_b = s_{11} - 2s_{01}$. As expected, this is the same dynamical law as for the strictly diploid model, in which case the dynamics of the selected allele are given by eq. (8) but with $s_a = s_{01}$. This enables us to identify a pair of 'effective' selection coefficients,

$$s_{01}^* = s_1 + s_{01}$$
$$s_{11}^* = 2s_1 + s_{11}, \tag{9}$$

so that, for weak selection, a diploid biallelic model with parameters $s_{01}^*$ and $s_{11}^*$ yields the same allele frequency dynamics[3] as a haplodiplontic model with parameters $s_1, s_{01}$ and $s_{11}$.

### S2.2. Equilibrium structure of the mainland-island model

We describe the equilibrium structure of the haplodiplontic single-locus deterministic mainland-island model for the biallelic case. The dynamics are given by the ODE

$$\dot{q} = -\dot{p} = m\Delta q + pq(s_a + s_b q) \tag{10}$$
$$= m\Delta q + pq(s_1 + s_{01} + (s_{11} - 2s_{01})q), \tag{11}$$

where $q$ is the frequency of the locally selected allele $A_1$, and $p = 1 - q$ is the frequency of the allele with relative fitness of 1 on the island when homozygous.

When $m = 0$ (no migration), there will be an admissible fixed point when either of the following conditions holds

$$s_{01} > -s_1 \text{ and } s_{01} > s_1 + s_{11} \tag{12}$$
$$s_{01} < -s_1 \text{ and } s_{01} < s_1 + s_{11}, \tag{13}$$

i.e. when there is *ploidally antagonistic selection*, diploid over- or underdominance, or both. The fixed point is obtained at

$$\tilde{p} = \frac{s_a + s_b}{s_a} = \frac{s_1 + s_{11} - s_{01}}{s_{11} - 2s_{01}} \tag{14}$$

This will correspond to a stable polymorphism whenever $s_{01} > 0$. This case was first analyzed in a discrete-time model by Scudo (1967).

[2] The same result can be obtained in a less cumbersome manner by first noting that, under the assumption of weak selection, allele frequency changes within a single alternation of generations are negligible, so that $w_{s,i}(p^*) = w_{s,i}(p)$ and $\bar{w}_s(p^*) = \bar{w}_s(p)$ in eq. (2).

[3] Note that a diploid model with these effective parameters does *not* yield the same mean fitness (and hence genetic load) as a haplodiplontic model in the original parameterization if we define mean fitness in the continuous time model as $\sum_i e^{s_i} p_i \big(\sum_j e^{s_{ij}} p_j\big)$.

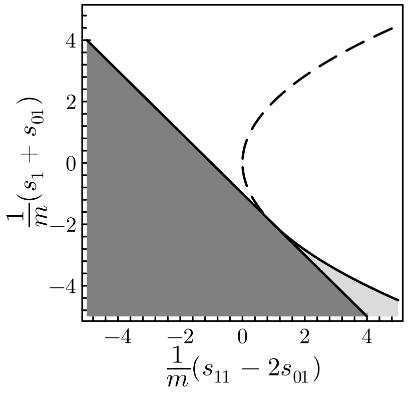


Fig. S26: Equilibrium and stability behavior of the single-locus biallelic haplodiplontic mainland-island model. The dark gray zone indicates the parameter region where there is a single protected stable polymorphic equilibrium. The light gray zone shows the parameter region where there is both a stable (but unprotected) and an unstable polymorphic equilibrium. $s_1$, $s_{01}$ and $s_{11}$ are the haploid and diploid selection coefficients for the invading allele ($A_1$) on the island.

Now consider $m > 0$. We shall assume that $\Delta q = q_{\text{mainland}} - q = 1 - q = p$, i.e. the mainland is fixed for the locally selected allele. To describe the equilibrium behavior, it is helpful to factor the dynamical law as

$$\dot{q} = mp\left(1 + \frac{s_a}{m}q + \frac{s_b}{m}q^2\right) \tag{15}$$

Linear stability at a fixed point $\tilde{q}$ is determined by

$$\frac{d\dot{q}}{dq}\Big|_{\tilde{q}} = (s_a - m) + 2(s_b - s_a)\tilde{q} - 3s_b\tilde{q}^2 \tag{16}$$

If $s_b = 0$, we have an effectively haploid model (i.e. *genic selection*), and will have a stable polymorphic equilibrium at $\tilde{q} = -m/s_a$ whenever $m < -s_a$, and a stable boundary equilibrium at $\tilde{q} = 1$ when $m > -s_a$. When $s_b \neq 0$, polymorphic equilibria, when they exist, will correspond to the roots of the quadratic expression in parentheses in eq. (15). These are

$$q_-, q_+ = \frac{-s_a/m \pm \sqrt{(s_a/m)^2 - 4s_b/m}}{2s_b/m}$$

We have the following biologically relevant equilibria:

i. $\tilde{q} = 1$ (*swamping*) is always a stable equilibrium when $m > -(s_a + s_b)$.
ii. When $0 < m < -(s_a + s_b)$ there is always a single stable polymorphic equilibrium at $q_-$ (dark gray zone in fig. S26) and $q_+$ will not lie in $[0, 1]$.
iii. When $-(s_a + s_b) < m < s_b$ and $4s_b/m < (s_a/m)^2 \iff 4m < s_a^2/s_b$, there is, besides the stable boundary equilibrium at $\tilde{q} = 1$, an unstable (repelling) equilibrium at $q_+$, and a stable polymorphic equilibrium at $q_-$ (light gray zone in fig. S26).

The relation between the key parameters $s_a/m$ and $s_b/m$ and the equilibrium behavior of the system when $m > 0$ is illustrated in fig. S26.

When condition (iii) holds, sharp thresholds for swamping are observed, in which case there is a certain critical allele frequency $p_c$ below which no local adaptation cannot be maintained whatever the migration rate. We can ask for which degree of dominance such sharp thresholds for swamping can possibly be observed. From eq. (15) we see that at an equilibrium which does not correspond to $p = 0$, the condition $f(q) = m + s_a q + s_b q^2 = 0$ holds. A sufficient condition for observing a sharp threshold is that $f$ obtains a maximum for some $q < 1$, hence that $f'(1) > 0$ where $f'(q) = s_a + 2s_b q$. This will be the case whenever $s_a + 2s_b > 0$. If we express this in terms of the effective selection and dominance coefficient so that $s_a = -s_e h_e$ and $s_b = -s_e(1 - 2h_e)$, this shows that critical behavior is expected as soon as $h_e > 2/3$.

### S2.3. Fixed point iteration algorithm

Our approximations yield a system of equations for the expected allele frequencies and heterozygosities on the island which are coupled through the gff. To calculate expected allele frequencies and allele frequency distributions at equilibrium, we solve the system self-consistently by performing a fixed point iteration. In words: for a given initial set of allele frequencies and heterozygosities, we calculate the gff at each locus using eq. (4); using these gff values, we next calculate expected allele frequencies and heterozygosities at each locus using numerical quadrature. This process is repeated until convergence. The algorithm is more formally outlined in algorithm S1.

**Algorithm S1** Fixed point iteration for calculating the expected allele frequency and expected heterozygosity on the island.

**Require:** Initialization $p^{(0)} = (p_1^{(0)}, \dots, p_L^{(0)})$, tolerance $\epsilon$
$(pq)^{(0)} \leftarrow (p_1^{(0)} q_1^{(0)}, \dots, p_L^{(0)} q_L^{(0)})$
$n \leftarrow 1, \Delta \leftarrow \infty$
**while** $\Delta > \epsilon$ **do**
  **for** $j = 1, \dots, L$ **do**
    $m_{e,j}^{(n)} \leftarrow \exp\left[\sum_{i \neq j} s_{i,a} q_i^{(n-1)} + s_{i,b}(p_i q_i)^{(n-1)}\right]$
    $p_j^{(n)} \leftarrow \int_0^1 p\phi(p; N_e, u, m_{e,j}^{(n)}, s_j)dp$
    $(p_j q_j)^{(n)} \leftarrow \int_0^1 p(1-p)\phi(p; N_e, u, m_{e,j}^{(n)}, s_j)dp$
  **end for**
  $\Delta \leftarrow \sum_j (p_j^{(n)} - p_j^{(n-1)})^2$
  $n \leftarrow n + 1$
**end while**
**return** $p^{(n)}, (pq)^{(n)}$

### S2.4. Swamping thresholds for the deterministic multilocus model

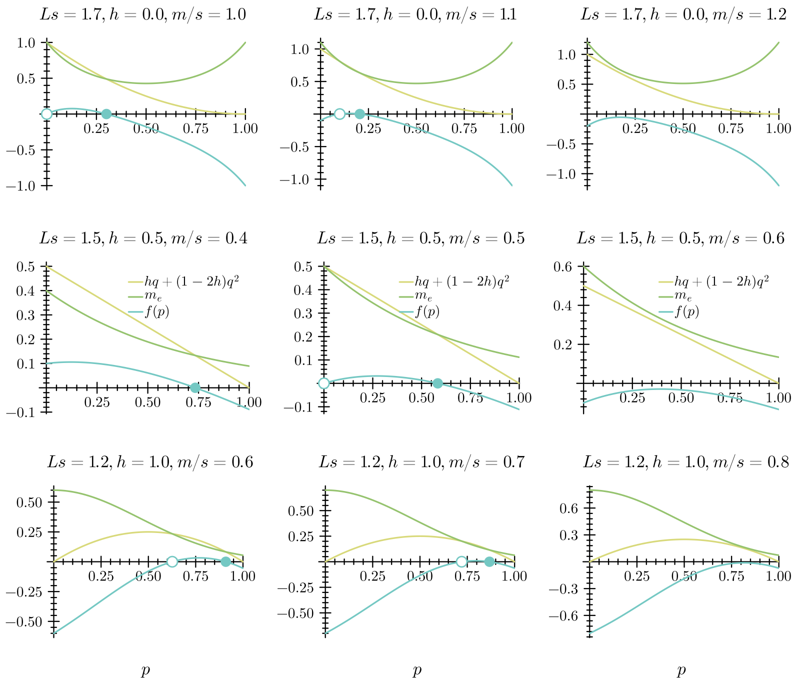


Fig. S27: Critical swamping thresholds for the multilocus model. Equilibria of the multilocus system correspond to the zeros of $f(p) = hq + (1-2h)q^2 - m_e/s$. Examples for $f(p)$ in the case with dominant local adaptation (top row), additive local adaptation (middle row) and recessive local adaptation (bottom row) near the critical point. The stable equilibrium is indicated by a filled dot, the unstable by an unfilled dot. When there is bistability, i.e. both a stable and unstable equilibrium, the critical migration rate at which the two equilibria collide and cease to exist corresponds to the value of $m$ for which $f(p)$ reaches its maximum in the critical point, so that both $f(p) = 0$ and $f'(p) = 0$ are satisfied.

We now take a closer look at the equilibria and critical behavior of the deterministic multilocus model with a homogeneous genetic architecture, expressed in terms of the effective selection and dominance coefficients $s$ and $h$ (eq. (10)). Clearly, $p = 0$ is always a solution of eq. (10), and it will correspond to a locally stable equilibrium (i.e. swamping) whenever $m/s > 1 - h$. Other equilibria, when they exist, are given by the zeros of the function

$$f(p) = hq + (1-2h)q^2 - \frac{m}{s} g[p] \tag{17}$$

Note that $g[p] > 0$ and we assume $s > 0$, so that for any fixed $h$, as $m$ increases, there will indeed be a critical migration rate beyond which $f(p) < 0$, from which point onwards the only stable equilibrium will be $p = 0$. At the critical point, the equilibrium allele frequency

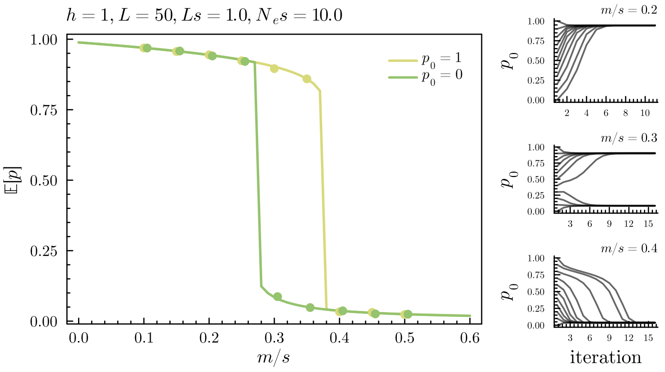


Fig. S28: Different apparent equilibria depending on initial conditions. In the left plot, the yellow line ($p_0 = 1$) indicates the expected allele frequencies as determined using the fixed point iteration of algorithm S1, starting with $p^{(0)} = (1, 1, \dots, 1)$ (i.e. secondary contact, maximal initial differentiation), whereas the green line assumes $p^{(0)} = (0, 0, \dots, 0)$ (no initial differentiation). The dots show results from individual-based simulations with the same initial conditions (50000 generation, keeping the last 25000 and subsampling every 5 generations). The three plots on the right show the evolution of the fixed point iteration for different initial initial conditions $p^{(0)} = (p_0, p_0, \dots, p_0)$ for three values of $m$. Note the bifurcation of the dynamical system defined by the algorithm: for $m/s = 0.2$ and $m/s = 0.4$ there is a single globally stable fixed point, whereas for $m/s = 0.3$, there are two locally stable fixed points.

will satisfy the additional constraint $f'(p) = 0$ (see fig. S27), i.e.

$$f'(p) = h + 2(1-2h)q - 2Lm(1 - 3h - 2(1-2h)q)g[p] = 0 \tag{18}$$

We can solve eq. (17) for $g[p]$, and then plug in $g[p]$ in eq. (18). This yields a cubic polynomial in $p$ which can be solved for the allele frequency $p_c$ at the critical point:

$$0 = Ls((1-h)^2 - 2p^3(1-2h)^2 + p^2(14h^2 - 17h + 5) - p(7h^2 - 11h + 4)) - 1 + \frac{3}{2}h + (1-2h)p \tag{19}$$

We can then plug $p_c$ into eq. (17) and solve for $m_c/s$. While the general expressions yield not much insight, we can focus on a number of special cases.

Firstly, in the additive case ($h = 0.5$), a critical point different from $1 - h$ appears when $Ls > 1$, in which case the equilibrium frequency at the critical point will be $p_c = 1 - 1/Ls$. The corresponding critical migration rate is

$$\frac{m_c}{s} = \frac{e^{Ls-1}}{2Ls}.$$

In the case where local adaptation is due to dominant alleles ($h = 0$), we have again critical behavior as soon as $Ls > 1$, with the swamping threshold occurring at $m/s = 1$ otherwise. In this case, we find

$$p_c = \frac{3}{4} - \frac{\sqrt{Ls(Ls+8)}}{4Ls} < \frac{1}{2}, \qquad \frac{m_c}{s} = \left(\frac{1}{4} + \frac{\sqrt{Ls\,(Ls+8)}}{4Ls}\right)^2 e^{\frac{Ls}{4} + \frac{\sqrt{Ls(Ls+8)}}{4} - 1}.$$

In contrast with the additive case (where as $Ls$ increases, arbitrary equilibrium differentiation can be maintained near the critical point), equilibrium differentiation will be below 0.5 near $m_c$ when $h = 0$. Lastly, for recessive local adaptation ($h = 1$), we have bistable critical behavior for all $Ls > 0$. The equilibrium frequency at the critical point is always larger than 1/2 and is given by the zeros of the cubic polynomial

$$4Lsp^3 - 4Lsp^2 + 2p - 1 = 0$$

for which we have no simple expressions. A fair approximation for $Ls < 1.5$ is given by

$$p_c \approx \frac{1}{2} + \frac{Ls}{4}, \qquad \frac{m_c}{s} \approx \left(\frac{1}{4} - \frac{(Ls)^2}{16}\right) e^{\left(\frac{Ls}{2}\right)^3 + \left(\frac{Ls}{\sqrt{2}}\right)^2 + \frac{Ls}{2}}$$

The swamping threshold is seen to increase strongly with increasing $Ls$.

### S2.5. Sensitivity to initial conditions

Although the equilibrium allele frequency distribution should be independent of the initial condition (the individual-based model can be thought of as an ergodic Markov chain on the space of $N$ $L$-locus genotypes), for appreciable $Ls$, the observed allele frequency distribution in any finite-time simulation can depend strongly on the initial conditions (that is, as $Ls$ increases, stochastic jumps between the different modes of the $L$-dimensional joint allele frequency distribution become increasingly less likely, and occur on time scales that are neither biologically relevant nor computationally feasible). This is especially true in the strongly recessive case ($h > 2/3$) and when LD is substantial. This is similar to the behavior in the deterministic model: when $Ls$ or $h$ is sufficiently large, and sharp swamping thresholds appear, the system is bistable, and the polymorphic equilibrium cannot be reached when the initial condition corresponds to a state of no or little differentiation.

The fixed point iteration will in that case converge to an expectation computed near one of the modes of the allele frequency distribution (see fig. S28 for an illustration). In other words, considering the fixed point iteration outlined in algorithm S1 as a discrete dynamical system, and treating $m/s$ as a bifurcation parameter, two bifurcation points occur succesively, as shown in fig. S28. For small $m/s$, a single globally stable polymorphic equilibrium is obtained. After the first bifurcation point, this equilibrium ceases to be globally stable, and a second locally stable equilibrium corresponding to almost no differentiation appears. After the second bifurcation point the lower equilibrium becomes globally stable. The region of parameter space where the two stable equilibria coexist corresponds to the situation where the assumption of population genetic equilibrium becomes questionable, where the state of the population after a large but finite time of evolution depends strongly on the detailed history of the population.

### S2.6. Distribution of fitness effects (DFE) models

#### S2.6.1. Independent selection and dominance coefficients

For the independent model, we assume, for $i = 1, \dots, L$,

$$s_i \sim \text{Gamma}(\kappa, \lambda)$$
$$h_i \sim \text{Beta}(\alpha, \beta),$$

where $\kappa$ is the shape parameter of the Gamma distribution, and $\lambda$ the rate parameter (i.e. the Gamma distribution with density $f(s) = \Gamma(\kappa)^{-1}\lambda^\kappa s^{\kappa-1}e^{-\lambda s}$). The mean is $\kappa/\lambda$, so that $\lambda = \kappa/\bar{s}$. The variance is $\kappa/\lambda^2 = \bar{s}^2/\kappa \propto \kappa^{-1}$ and the excess kurtosis is similarly $\propto \kappa^{-1}$. Decreasing $\kappa$ therefore increases simultaneously the variance and kurtosis of $s$ across the barrier. For $\kappa = 1$, this reduces to the Exponential distribution with rate $\lambda$. See fig. S29 (A) and fig. S30 for examples.

#### S2.6.2. Logistic model

In the logistic model, we assume, for $i = 1, \dots, L$,

$$s_i \sim \text{Gamma}(\kappa, \lambda)$$
$$\text{logit}\, h_i | s_i \sim \text{Normal}(a + b \log s_i, \sigma^2),$$

where $\text{logit}\, h = \log \frac{h}{1-h}$ is the logit transform. In other words, we assume $h$ to be distributed according to a linear regression on $\log s$ with slope $a$ and intercept $b$, on a logit scale. The marginal density of $h$ is then

$$f(h) = \frac{1}{h(1-h)} \int_0^\infty \text{N}[\text{logit}\, h; a + b\log(s), \sigma] \text{G}(s; \kappa, \lambda) ds \ ,$$

where $\text{N}(\cdot; \mu, \sigma)$ and $\text{G}(\cdot; \kappa, \lambda)$ denote the density functions for the Normal distribution (with mean $\mu$ and standard deviation $\sigma$) and Gamma distribution respectively. Instead of setting the $a$ and $b$ parameters directly, we parameterize the regression by choosing two reference pointst, $s^{(1)}$ and $s^{(2)}$, together with their respective expected dominance coefficients $h^{(1)} = \mathbb{E}[h|s^{(1)}]$ and $h^{(2)} = \mathbb{E}[h|s^{(2)}]$ using

$$b = \frac{\text{logit}\, h^{(2)} - \text{logit}\, h^{(1)}}{\log s^{(2)} - \log s^{(1)}}$$
$$a = \text{logit}\, h^{(1)} - b \log s^{(1)}$$

This model is also illustrated in fig. S29 (B) and fig. S30.

#### S2.6.3. Model after Caballero and Keightley (1994)

In the model of Caballero and Keightley (1994) (see also Zhang *et al.* (2004) and discussion in Agrawal and Whitlock (2011)), referred to as CK94, we assume

$$s_i \sim \text{Gamma}(\kappa, \lambda)$$
$$h_i^* | s_i \sim \text{Uniform}(0, e^{-Ks})$$
$$h_i = 1 - h_i^*$$

It should be noted that this distribution is supposed to be a reasonable model for dominance coefficients of deleterious mutations at mutation-stabilizing selection equilibrium, where mutations of large effect segregating at appreciable frequencies tend to be recessive. In

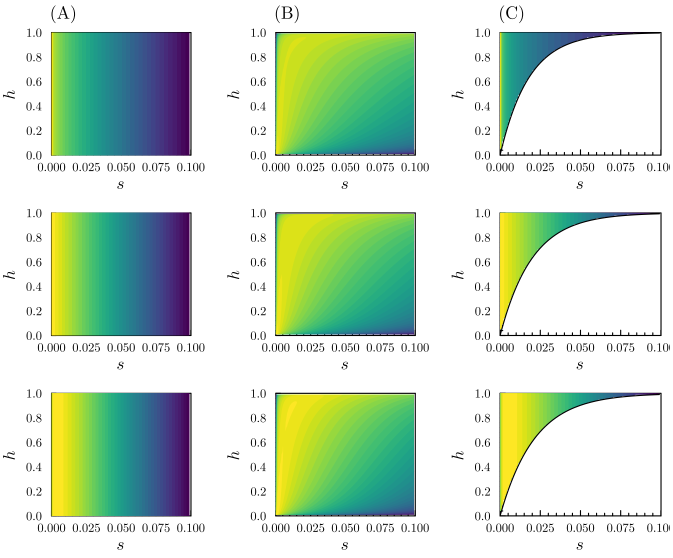


Fig. S29: Joint distributions for an example of each of the three DFE models outlined in section S2.6. The joint probability density is shown on a logarithmic scale, with yellow marking high density and (deep) blue low density. The marginal density for the selection coefficient is a Gamma distribution with $\kappa = 2/3, 1, 3/2$ in the top, middle and bottom row respectively (see section S2.6 for the relevant definitions). (A) Independent selection and dominance coefficients. (B) The logistic model with $a = 9.2$ and $b = 2$. (C) The CK94 model with $\bar{h} = 1/3$.

our case, we regard this as the distribution of dominance coefficients of *mutant* alleles on the *mainland* that constitutes the standing variation which is the source of locally adaptive alleles during the initial polygenic response (which we do not explicitly model). These are hence the dominance coefficients of the locally *beneficial* alleles on the *island* (assuming dominance coefficients to be constant across environments and genetic backgrounds). The $h_i$ as we defined them are however the dominance coefficients of the invading wild-type alleles from the mainland over the locally beneficial ones, so that we use $h_i = 1 - h_i^*$ where the $h_i^*$ are distributed according to the CK94 model. We set the $K$ parameter so that $\mathbb{E}[h^*] = \bar{h}$ for some $\bar{h}$, i.e.

$$K = \lambda\left((2\bar{h})^{-\frac{1}{\kappa}} - 1\right).$$

The marginal density for $h^*$ is

$$\begin{aligned}
f(h) &= \int_0^{-\frac{\log h}{K}} \frac{\lambda^\kappa}{\Gamma(\kappa)} \lambda^\kappa s^{\kappa-1} e^{-(\lambda-K)s} ds \\
&= \left(\frac{\lambda}{\lambda-K}\right)^\kappa \int_0^{-\frac{\log h}{K}} \frac{(\lambda-K)^\kappa}{\Gamma(\kappa)} \lambda^\kappa s^{\kappa-1} e^{-(\lambda-K)s} ds = \left(\frac{\lambda}{\lambda-K}\right)^\kappa \int_0^{-\frac{\log h}{K}} \mathrm{G}(s;\kappa,\lambda-K) ds \\
&= \left(\frac{\lambda}{\lambda-K}\right)^\kappa \frac{\gamma\left(\kappa, -(\lambda-K)\frac{\log h}{K}\right)}{\Gamma(\kappa)}
\end{aligned}$$

where $\gamma$ is the lower incomplete gamma function. This model is also illustrated in fig. S29 (C) and fig. S30. To study the effect of a negative correlation between $s$ and $h$ (i.e. where strongly selected locally beneficial alleles tend to be dominant), we use the same model, but with $h_i = h_i^*$. We refer to this model as CK94* (see fig. S22).

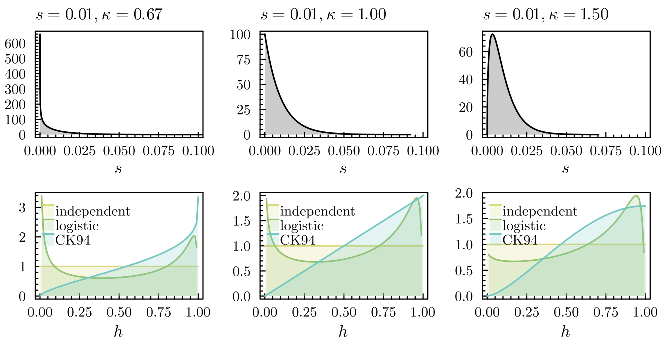


Fig. S30: Marginal distributions of $s$ and $h$ for the three example DFE model shown in fig. S29.
